# Shared heritability of face and brain shape distinct from cognitive traits

**DOI:** 10.1101/2020.08.29.269258

**Authors:** Sahin Naqvi, Yoeri Sleyp, Hanne Hoskens, Karlijne Indencleef, Jeffrey P. Spence, Rose Bruffaerts, Ahmed Radwan, Ryan J. Eller, Stephen Richmond, Mark D. Shriver, John R. Shaffer, Seth M. Weinberg, Susan Walsh, James Thompson, Jonathan K. Pritchard, Stefan Sunaert, Hilde Peeters, Joanna Wysocka, Peter Claes

**Affiliations:** Department of Chemical and Systems Biology, Stanford University School of Medicine, Stanford, California, United States of America; Departments of Genetics and Biology, Stanford University School of Medicine, Stanford, California, United States of America; Department of Human Genetics, KU Leuven, Leuven, Belgium; Medical Imaging Research Center, University Hospitals Leuven, Leuven, Belgium; Department of Electrical Engineering, ESAT/PSI, KU Leuven, Leuven, Belgium; Department of Neurosciences, KU Leuven, Leuven, Belgium; Neurology Dept, University Hospitals Leuven, Leuven, Belgium; Department of Imaging and Pathology, Translational MRI, KU Leuven, Leuven, Belgium; Department of Biology, Indiana University Purdue University Indianapolis, Indianapolis, Indiana, United States of America; Applied Clinical Research and Public Health, School of Dentistry, Cardiff University, Cardiff, United Kingdom; Department of Anthropology, Pennsylvania State University, State College, Pennsylvania, United States of America; Department of Human Genetics, University of Pittsburgh, Pittsburgh, Pennsylvania, United States of America; Department of Oral Biology, Center for Craniofacial and Dental Genetics, University of Pittsburgh, Pittsburgh, Pennsylvania, United States of America; Department of Anthropology, University of Pittsburgh, Pittsburgh, Pennsylvania, United States of America; Department of Psychology George Mason University, Fairfax, Virginia, United States of America; Department of Developmental Biology, Stanford University School of Medicine, Stanford, California, United States of America; Howard Hughes Medical Institute, Stanford University School of Medicine, Stanford, California, United States of America; Murdoch Children’s Research Institute, Melbourne, Victoria, Australia

**Author notes:** Joint first authorship (Sahin Naqvi, Yoeri Sleyp). Joint last authorship (Joanna Wysocka, Peter Claes).

## Abstract

Evidence from both model organisms and clinical genetics suggests close coordination between the developing brain and face^1–8^, but it remains unknown whether this developmental link extends to genetic variation that drives normal-range diversity of face and brain shape. Here, we performed a multivariate genome-wide association study of cortical surface morphology in 19,644 European-ancestry individuals and identified 472 genomic loci influencing brain shape at multiple levels. We discovered a substantial overlap of these brain shape association signals with those linked to facial shape variation, with 76 common to both. These shared loci include transcription factors with cell-intrinsic roles in craniofacial development, as well as members of signaling pathways involved in brain-face crosstalk. Brain shape heritability is equivalently enriched near regulatory regions active in either brain organoids or in facial progenitor cells. However, brain shape association signals shared with face shape are distinct from those shared with behavioral-cognitive traits or neuropsychiatric disorder risk. Together, we uncover common genetic variants and candidate molecular players underlying brain-face interactions. We propose that early in embryogenesis, the face and the brain mutually shape each other through a combination of structural effects and paracrine signaling, but this interplay may have little impact on later brain development associated with cognitive function.

## MAIN

The human cerebral cortex forms the outer layer of gray matter of the brain and underpins cognitive function. It is characterized by complex folding patterns that vary between species and individuals^9,10^. Family- and twin-based studies indicate substantial heritability of brain shape^11,12^, and a recent genome-wide association study (GWAS) found that brain shape is highly polygenic and shows genetic correlations with a broad range of neuropsychiatric disorders and behavioral-cognitive phenotypes^13^. These studies focused on pre-defined, univariate measures of brain shape, such as total or regional surface area, extracted from structural magnetic resonance imaging (MRI) scans^14^, and which cannot capture the morphological complexities of the cortical surface. We recently developed a data-driven approach to phenotyping complex, multidimensional traits^15^; this fully multivariate approach, when applied to facial surface images, revealed a large number of novel loci associated with variation in human face shape^15,16^. Here, we implemented this approach to discover associations between common genetic variants and brain shape, using MRI data from largely healthy, middle-aged participants in the UK Biobank (UKB).

In addition to sharing complex morphologies, the development of the brain and face is highly integrated as a result of shared developmental lineage, spatial proximity, and signaling crosstalk between the two structures. Early in embryonic development, the rostral end of the ectodermally-derived neural tube gives rise to the forebrain, which in turn gives rise to the cerebrum that encompasses the cerebral cortex. Just before forebrain formation, a subset of neuroepithelial cells within the neural folds give rise to facial progenitor cells called cranial neural crest cells (CNCCs). Following specification, CNCCs undergo an epithelial-to-mesenchymal (EMT) transition and migrate ventrally, later giving rise to most of the craniofacial skeleton and connective tissue. Early growth rates of the brain can modulate both the positioning and outgrowth of the facial prominences^1,2^, as well as induce flexion and bone deposition of the CNCC-derived basicranial bones^3,17^ and neurocranial sutures^18,19^, respectively. Finally, paracrine factors secreted by either the developing forebrain^20–23^ or CNCCs^5,6,24^ modulate the development of the face or brain, respectively.

These physical and molecular interactions have been detailed by studies in the developing chick and mouse embryos, but are also supported by widespread co-occurrence of neurodevelopmental and craniofacial malformations in rare human syndromes^7^. This phenomenon was noticed as early as 1964, when Demyer *et al*. coined the phrase “the face predicts the brain” to describe the correlation between the severity of brain abnormalities and facial malformations in patients with holoprosencephaly^8^. While in some cases this co-occurrence may be caused by pleiotropic functions of the affected gene, a number of such human syndromes have been mapped to genes known to function in brain-face crosstalk through paracrine signaling^25–27^. Nonetheless, close developmental links between face and brain are often underappreciated; whether and how they extend to common human genetic variation that influences the diversity of brain and face shape is unknown.

### Multivariate GWAS of brain shape

We adapted our previously published data-driven phenotyping approach^15^ to brain shape, as measured by MRI scans of 19,644 individuals in UKB. Participants included were of primarily European ancestry, such that results do not pertain to cross-population differences in brain shape. All of our analyses focused on the mid-cortical surface (midway between the white-grey matter interface and the pial surface with the cerebrospinal fluid, as extracted using FreeSurfer^14^), which we refer to as brain shape. Given the complete dataset of mid-cortical surfaces each represented by a homologous mesh of spatially dense 3D vertices, the method segments brain shape in a global-to-local manner, yielding multivariate brain segments at different hierarchical levels of scale. Within each segment, principal component analysis (PCA) is used to describe effects in multivariate shape-space explaining between-individual variation, and canonical correlation analysis (CCA) is used to define, for each variant tested in the genome, the linear combination of PCs maximally associated with single nucleotide polymorphism (SNP) dosage. In agreement with findings of nonzero but low heritability of thickness and surface area asymmetry^28^, we observed that independent processing and GWAS of left and right hemispheres showed highly concordant results (Extended Data Fig. 1). Therefore, all subsequent analyses were performed using the left-right hemisphere averaged surface data.

Applying this pipeline to the UKB MRI data, we defined 285 hierarchical segments (Fig. 1, Supplementary Table 1), decomposing brain shape into different levels of detail, from larger brain segments with more integrated shape variation, to more smaller brain segments with more local effects. Each hierarchical level is a bipartition of its parent; thus, the first level consisted of the entire brain, while the second and third levels segmented the whole brain into halves and quadrants, respectively, and the final, ninth level resulted in numerous smaller segments (Figure 1b, right). Many smaller segments from the seventh hierarchical level onwards were discarded due to their small surface areas, resulting in fewer total segments than the 511 (2^9^ - 1) expected. The segmentation broadly agreed with the commonly-used Desikan-Killiany^29^, Destrieux^30^, and Glasser^31^ atlases of brain regions (Extended Data Fig. 2). In total, we conducted 285 multivariate GWAS using CCA, each corresponding to one segment. 38,630 SNPs showed genome-wide significant (*P* < 5 x 10^-8^) association with brain shape variation in at least one segment; of these, 23,413 reached study-wide significance (*P* < 2.07 x 10^-10^ as assessed by permutation, see Methods) in at least one segment. Collapsing these SNPs into independent signals based on linkage disequilibrium and distance yielded 472 and 242 loci reaching genome- and study-wide significance, respectively (Supplementary Table 2). Most of the 472 loci showed effects on multiple segments (305/472, 65%), and many showed effects on multiple quadrants (158/472, 33%) (Figure 1, Supplementary Table 2), consistent with global-to-local effects at multiple levels of brain shape. Associations between these loci and brain shape were generally depleted from the frontal lobe segments (except for the most anterior orbitofrontal cortex) and enriched in the occipital and temporal lobe segments (Extended Data Fig. 3), mostly in agreement with point-wise heritability estimates across the brain surface (Extended Data Fig. 4).

**Fig. 1:**
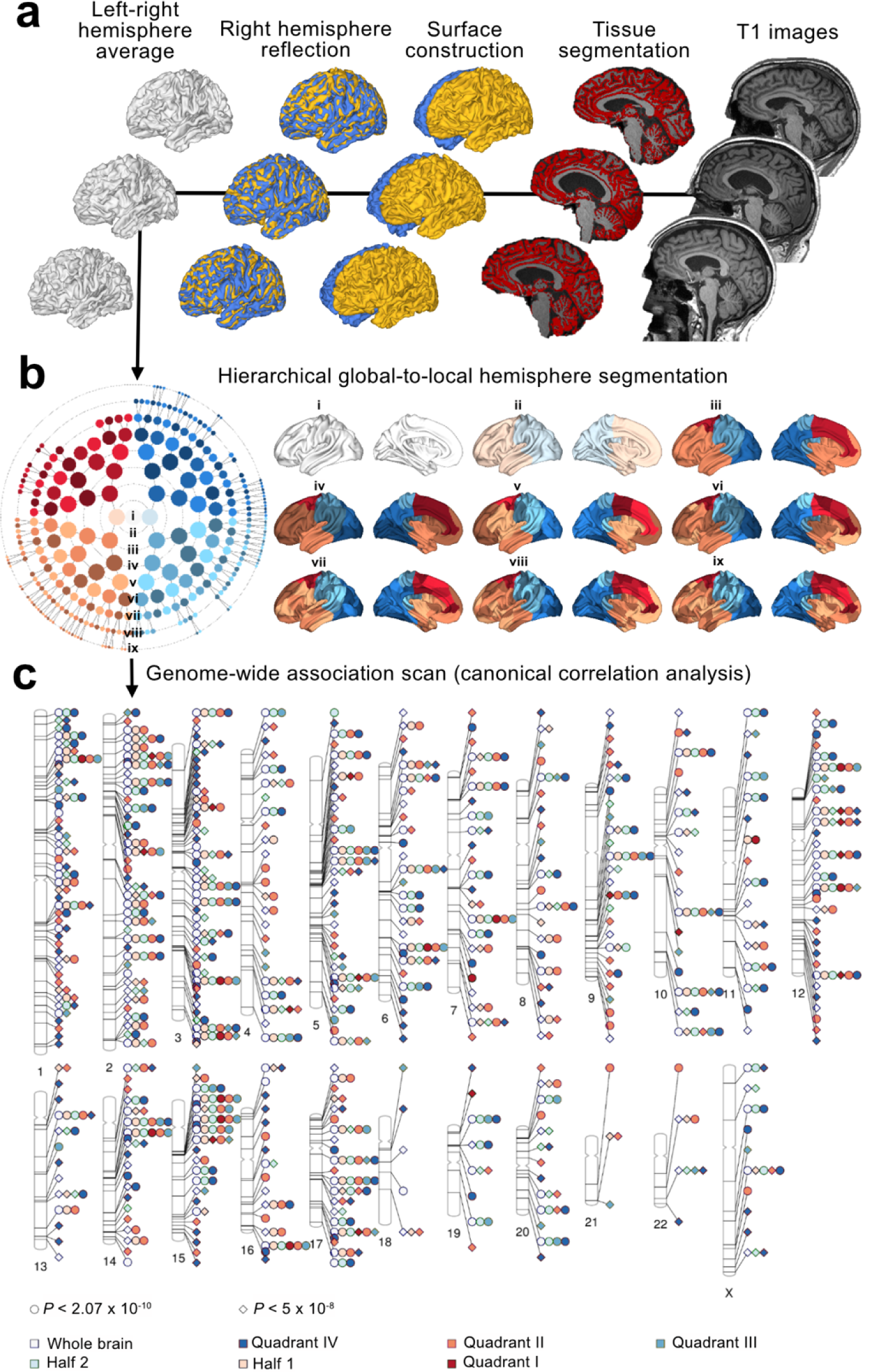
Multivariate genome-wide association study (GWAS) of brain shape. **a**, Upstream processing of UKB MRI images. **b**, in the polar dendrogram on the left, each concentric ring of filled circles corresponds to a hierarchical level (labeled i-ix) shown on the right, and the filled circle colors correspond to the respective segments in the same hierarchical level. **c**, Ideogram showing genomic locations and regional effects of 472 genome-wide significant loci for brain shape. Circles and diamonds represent associations passing the study-wide or genome-wide significance thresholds, colors represent broad regions of the brain with the indicated effects.

We assessed the overlap between the 472 loci and previous GWAS of brain surface areas or subcortical volumes^13,32–36^. The 472 loci recapitulated 27-78% of the associations reported in previous studies; the highest overlap of 78% was with a recent study of univariate brain surface area^13^, which is the phenotype most comparable to the shape measures studied here (Extended Data Table 1). In total, of the 472 loci, 121 overlapped with those reported in previous studies on brain surface area or subcortical volume, while 351 represent novel associations with brain morphology. To assess the reproducibility of the 472 loci on the same shape measures, we analyzed MRI data from the Adolescent Brain Cognitive Development (ABCD) study^37^. Of the 472 loci, 466 were available for replication testing (see Methods). At 5% FDR, we replicated at least one associated segment for 305 of 466 (65.4%) loci, and 2,645 of 3,586 (73.8%) tested locus-segment combinations (Supplementary Table 3). This replication rate is notable given the substantial difference in age of the ABCD cohort (9-10 years versus 40-70 years in UKB). Thus, despite the known continued growth and morphological changes of the brain throughout adolescence and into adulthood^38^, the high reproducibility of GWAS results between the two cohorts suggests that many of the observed associations with brain shape originate during development and are maintained throughout life.

We next used FUMA^39^ and GREAT^40^ to identify pathways enriched among genes near the 472 loci, as well as curated gene panels used to guide rare disease diagnoses from whole-genome sequencing^41^ to identify disease associations (see Methods for details). As expected, we found strong enrichment for brain-specific processes (i.e. neurogenesis, axonogenesis, neuron differentiation, nervous system development, neuron projection guidance), morphogenesis-related processes (i.e. anatomical structure morphogenesis, animal organ morphogenesis), and neurodevelopmental disorders (i.e. intellectual disability, malformations of cortical development, ciliopathies). We also observed a weak enrichment of terms related to the formation and closure of the neural tube, suggesting that early developmental events impact adult brain shape variation. Surprisingly, we also observed strong enrichment of terms related specifically to CNCC development and migration, as well as weaker enrichments in broader terms encompassing skeletal system development, chondrogenesis, and osteogenesis (Supplementary Table 4). Furthermore, strong and weak enrichments were also found for craniosynostosis and clefting gene panels, respectively. These enrichments suggested a link between variation in brain shape and craniofacial skeletal development, which we set out to explore further.

### Loci affecting both brain and face shape

To more directly test for sharing of genetic effects between brain and face shape, we intersected the 472 loci described in this study with 203 loci we previously associated with face shape variation in European-ancestry individuals through a similar, open-ended phenotyping approach^16^. Thirty-seven of the loci for brain shape are in linkage (r^2^ > 0.2) with at least one of the face shape loci, significantly above random expectation (*P* = 2.03 x 10^-22^, OR = 10.6) and greater than the overlap with other traits that have similar numbers of genome-wide significant associations in the NCBI-EBI GWAS Catalog^42^ (Extended Data Fig. 5). Identifying signals showing genome-wide significant association with one of brain or face shape and genome-wide suggestive (*P* < 5 ⨯ 10^-7^) association with the other resulted in 76 brain-face shared loci (Figure 2a), which we carried forward for further analysis.

**Fig. 2:**
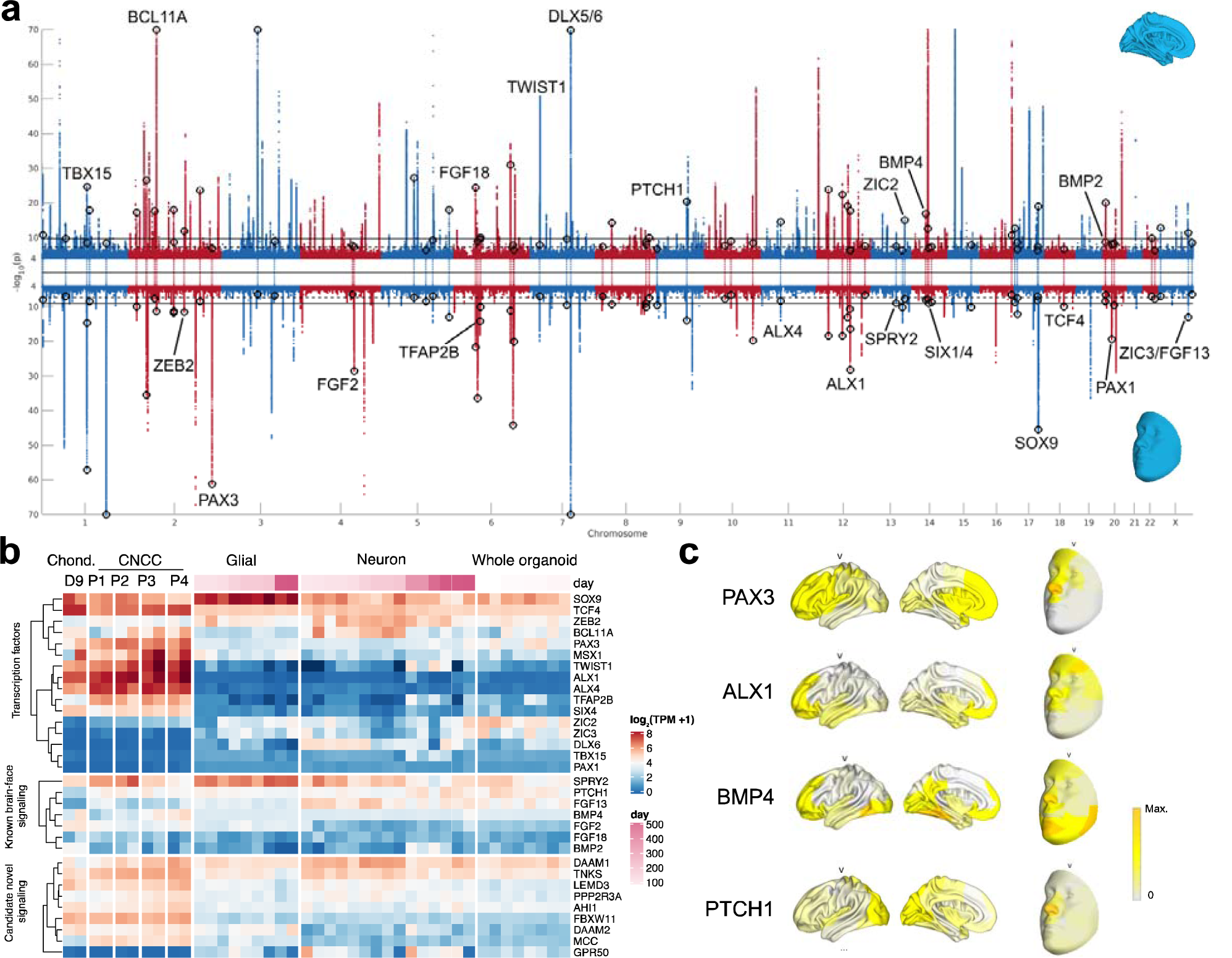
Loci affecting both brain and face shape. **a**, Miami plot of GWAS for brain (top) and face (bottom) shape. For each SNP, p-values aggregated across all brain or face segments are plotted. All 76 loci reaching genome-wide (*P* < 5 ⨯ 10^-8^) significance in one study and genome-wide suggestive (*P* < 5 x 10^-7^) significance in the other are highlighted by unfilled circles. Loci near candidate genes highlighted in the text and in **b** and **c** are labeled, generally on the side where they show greater significance of association. **b**, Expression (in transcripts per million, TPM) of candidate genes near brain-face shared loci in cranial neural crest cells (CNCCs) of different passages, representing different stages of maturation, from early (P1) to late (P4) and their chondrocyte (Chond. D9) derivatives^49^ (left), and three dimensional forebrain organoids at various stages of differentiation^50^ (right), further sorted into glial or neuronal lineages or profiled as whole organoids. **c**, Regional phenotypic effects of four candidate loci, showing effects of linked SNPs on brain (left) or face (right) shape. Segments shown are of hierarchical level v, - log10(p-values) are normalized to the maximum at each locus. Full face and brain images from all 76 brain-face shared loci corresponding to all hierarchical levels can be found online (see Data Availability)

Genes near the 76 brain-face shared loci were strongly enriched for disease associations, including “skeletal disorders” and “hearing and ear disorders”, consistent with the contribution of CNCCs to craniofacial skeleton and ear structures. We next scanned the 76 brain-face shared loci for candidate genes with known roles in craniofacial or brain development from human syndromes and/or mouse knockouts (Supplementary Table 5). We observed that many of the shared brain-face loci are associated with genes encoding transcription factors (TFs) involved in neural crest formation and/or craniofacial skeletal development. Some of those TFs (for example DLX5/6, SOX9, ZEB2, ZIC2, ZIC3, TCF4) have known functions in both neural crest and brain development, and this pleiotropy may account for the shared genetic signals observed between the face and the brain. However, other shared brain-face signals are associated with TFs thought to function primarily during neural crest rather than brain development, and whose mutations cause specific craniofacial defects; those TFs include ALX1 and ALX4 (associated with frontonasal dysplasias^43,44^), TWIST1 (associated with Saethre-Chotzen Syndrome^45,46^), PAX3 (associated with Waardenburg syndrome^47^), and TFAP2B (associated with CHAR syndrome^48^). Consistent with the primary role of these TFs in facial development, transcriptome analysis showed their high expression in in-vitro derived human CNCCs and their chondrocyte derivatives^49^, but low/no expression in either glia or neurons of human forebrain organoids spanning a wide range of developmental stages^50^ (Figure 2b). These observations suggest that genetic variants influencing regulation of key craniofacial TFs have a greater than previously appreciated impact on brain shape.

Interactions between face and brain can be architectural in nature, with the forebrain acting as a structural support for facial development, and facial skeletal structures flexing to accommodate early brain growth^4^. However, these interactions can also involve paracrine signaling, with fibroblast growth factor (FGF), Hedgehog, and bone morphogenetic protein (BMP) pathways having documented roles in mediating the signaling from the developing brain to the face^20–22^. Interestingly, genes encoding members of all three aforementioned pathways, FGF (*FGF2, FGF13, FGF18, SPRY2*), Hedgehog (*PTCH1*), and BMP (*BMP2, BMP4*) are among the shared brain-face association loci. For example, mutations in *PTCH1*, the receptor for the sonic hedgehog ligand, cause holoprosencephaly^51^, a congenital, structural forebrain anomaly also associated with a range of craniofacial malformations. Conversely, CNCCs secrete anti-BMP signaling molecules which modulate forebrain development^5,6^, and expression of these BMP antagonists is dependent on the SIX family TFs, whose perturbation in CNCCs is associated with craniofacial malformations, but also causes secondary pre-otic brain defects^52^; *SIX1/4* is also among the shared brain-face loci identified in our study (Figure 2a). We further note that a number of genes linked to regulation of other signaling pathways, for which prominent roles in brain-face crosstalk have yet to be described, including Wnt (*DAAM1, DAAM2, TNKS, AHI1, FBXW11, MCC*) and transforming growth factor beta (*LEMD3, PPP2R2A*) are among the shared brain-face association loci, suggesting new candidate genes and pathways for future functional exploration. Not unexpectedly, and in contrast to the craniofacial TFs, the signaling pathway ligands, receptors and regulators are variably expressed between the in-vitro derived human CNCCs and brain organoids (Figure 2b).

Phenotypically, these highlighted loci largely affect brain shape in the frontal and temporal lobes, and face shape in the forehead and nose, as exemplified by *PAX3* and *ALX1* (Figure 2c), consistent with the physical proximity of the frontonasal prominence and the forebrain during development. Phenotypic effects distinct from this pattern include effects of variants near *BMP4* and *DLX6* on jaw and chin morphology, consistent with their known roles in mandibular development^53,54^, and effects of variants near *PTCH1* on occipital lobe morphology (Figure 2c). Together, these results suggest that both cell-intrinsic mechanisms and paracrine signaling pathways contribute to the substantial number of loci with shared associations with brain and face shape.

### Genome-wide sharing of signals with neuropsychiatric disorders and behavioral-cognitive traits

We next asked whether the brain-face overlap among genome-wide significant loci was true across the genome, also considering GWAS of neuropsychiatric disorders and behavioral-cognitive traits. Genetic correlations between univariate traits can be computed from signed summary statistics using LD score regression (LDSC)^55^. However, this approach is not applicable to the unsigned statistics yielded by CCA. We therefore applied an alternative method of assessing genome-wide sharing of signals between two GWAS, summarizing SNP p-values within approximately independent LD blocks and computing Spearman correlations between the two summarized profiles (see Methods for details). When applied to pairs of univariate GWAS, the Spearman correlation method was largely concordant with, albeit generally smaller in magnitude than, unsigned estimates of genetic correlations by LDSC (Extended Data Fig. 6), indicating that it is a conservative, robust measure for quantifying genome-wide sharing of GWAS signals.

We first assessed sharing of association signals between 63 face segments and 285 brain segments (Supplementary Table 6). All four main facial quadrants, representing shape variation within the forehead, nose, lower face (mandible and cheeks) and philtrum, respectively, showed the most sharing with brain segments in the frontal lobe, particularly the most anterior portions such as the rostral prefrontal cortex, and the least sharing with segments in the parietal lobe (Figure 3a). Furthermore, among the facial quadrants, the forehead and nose showed more sharing with frontal lobe effects than the philtrum and lower face. These genome-wide correlations are consistent with the phenotypic effects of top brain-face shared loci (Figure 2c, Extended Data Fig. 7).

**Fig. 3:**
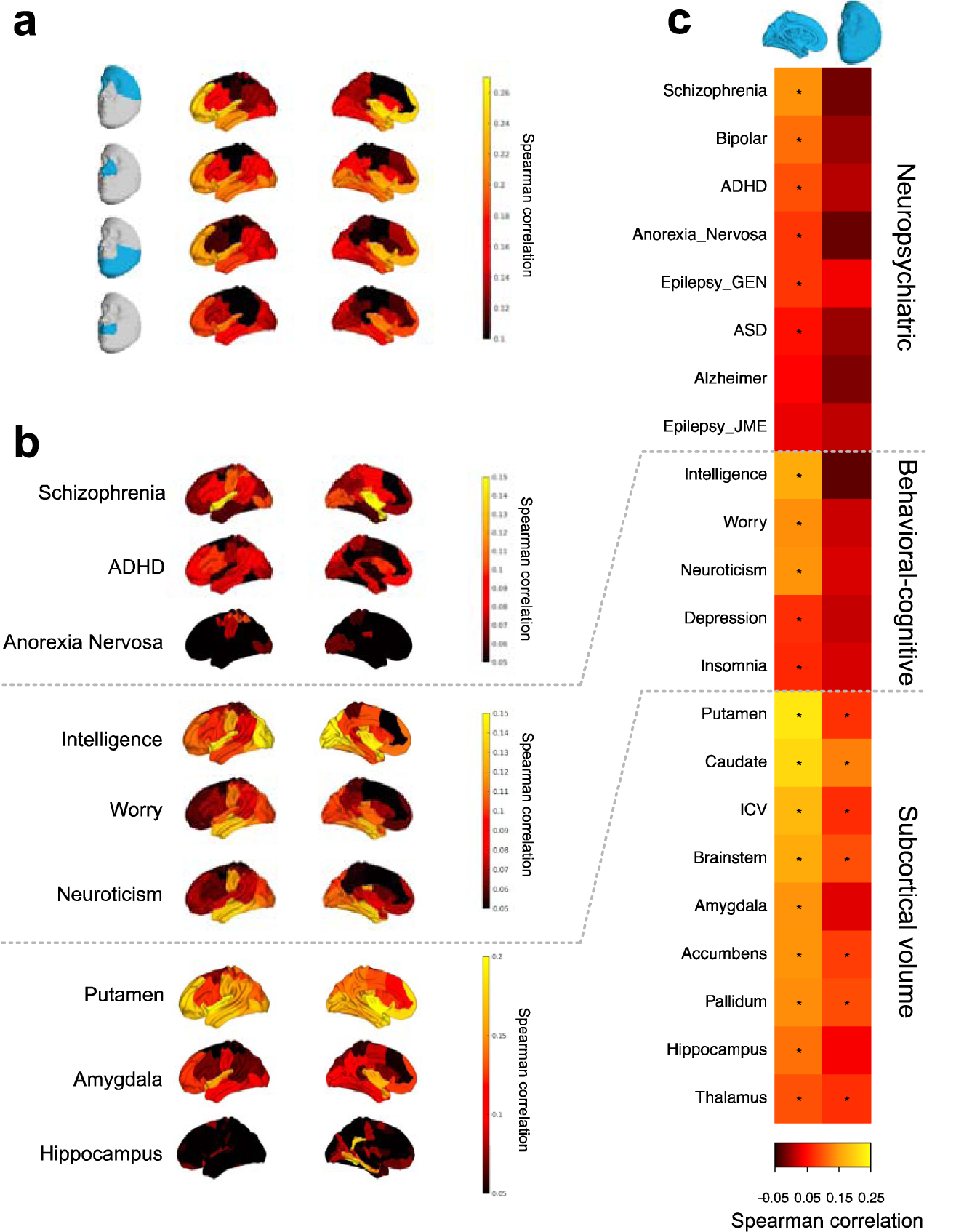
Genome-wide sharing of signals with neuropsychiatric disorders and behavioral-cognitive traits. Genome-wide sharing of signals between any two given GWAS was assessed by Spearman correlation of LD block-average SNP -log10(p-values) (Methods). **a**, Spearman correlations between GWAS of indicated facial quadrants and brain segments. **b**, Spearman correlations between GWAS of selected neuropsychiatric disorders, behavioral-cognitive traits, or subcortical volume measures and brain segments. All brain segments in **a**,**b** are from hierarchical level v segmentation, with the exception of Hippocampus, where hierarchical level vi segmentation shows a strong correlation of shape of the hippocampal region with volume. **c**, Spearman correlations between shape effects on the full brain (left) or face (right) with the indicated traits. * 5% FDR based on bootstrapped p-value (Methods). Images of brain-trait correlations at all six hierarchical levels can be found online (see Data Availability). Abbreviations: ADHD, Attention Deficity Hyperactivity Disorder ;GEN, generalized epilepsy; JME, juvenile myoclonic epilepsy; ICV, intracranial volume.

We next assessed sharing of signals with other phenotypes relevant to brain shape. We used publicly-available genome-wide summary statistics for a range of neuropsychiatric disorders, behavioral-cognitive traits, and subcortical brain volume measures from studies other than the UK Biobank, since our Spearman correlation measure does not control for sample overlap (Supplementary Table 7). Subcortical volume measures showed the most sharing with brain shape in the corresponding regions, but the magnitude of these correlations was relatively low (on par with sharing between brain and face shape), indicating that our multivariate GWAS approach detects many effects beyond those resulting from changes in relative subcortical volume (Figure 3b). We found that disorders with primarily developmental etiology and that manifest early in life showed substantial sharing with brain shape in regions previously linked to these disorders. For instance, schizophrenia and attention deficit hyperactivity disorder (ADHD) showed sharing with shape variation in the primary auditory^56,57^ and prefrontal cortex regions^58^, respectively. In contrast, we did not observe this association with cortical surface shape in Alzheimer’s disease, caused by plaque buildup and neurodegeneration much later in life. Associations with behavioral-cognitive traits such as intelligence, neuroticism and worry showed broader patterns of sharing with brain shape across multiple regions, reflecting the presumed involvement of distributed cortical regions in these traits^59–61^ (Figure 3b).

Finally, we compared the degree to face shape shares signals with neuropsychiatric disorders, behavioral-cognitive traits, and subcortical volume measures. Brain shape shares significant (FDR 5%) signal with most neuropsychiatric traits, as well as all behavioral-cognitive and subcortical volume traits analyzed. In contrast, face shape does not show significant sharing with any of the neuropsychiatric disorders or behavioral-cognitive traits, and significant but weaker sharing with the subcortical volume measures (Figure 3c). To confirm these patterns of sharing using standard univariate approaches for face shape, we performed GWAS on the most heritable individual PCs of full brain or face shape and computed standard genetic correlations using LDSC. Although genetic correlation estimates were noisy due to low heritability of shape GWAS using univariate approaches, they generally agreed with our Spearman correlation measure, finding non-zero genetic correlations between both brain and face shape and subcortical volume measures, and between brain shape and autism spectrum disorder (Extended Data Fig. 8). Thus, the substantial sharing of signals between brain and face shape (Figure 3a) is mostly independent of neuropsychiatric disorder risk and behavioral-cognitive traits, likely due to the fact that mutual influences of face and brain shape on each other involve phenotypic effects on brain shape distinct from those important for risk of neuropsychiatric disorders and behavioral-cognitive traits.

### Cell-types influencing brain and face shape

Our results thus far suggest that a substantial fraction of brain shape variation is underpinned by face shape variation, but that these effects are largely independent of effects shared between brain shape and other cognitive traits. To systematically test this idea further, we sought to identify the cell-types and tissues most enriched for heritability of brain shape, face shape, and other traits relevant to cognitive function. Partitioning heritability into cell-type specific functional annotations (i.e. open chromatin, enhancers, and promoters) via stratified LD score regression (S-LDSC) can prioritize trait-relevant cell-types and tissues, but was developed for univariate traits^62^; we thus sought to extend the theoretical framework of S-LDSC to multivariate traits such as the brain and face shape GWAS in this study. We proved that when applying unstratified LDSC^55^ to χ^2^ statistics obtained from multivariate traits with independent dimensions and further corrected for trait dimensionality, the LDSC-estimated heritability is equal to the average heritability of the component univariate traits (see Methods, Supplementary Note), a proof that we validated through separate LDSC heritability analysis of each PC making up the full face (Extended Data Figure 9). By extension, heritability enrichments obtained by applying S-LDSC on multivariate, corrected χ^2^ statistics partitioned by a given functional annotation represent the average heritability enrichment for each component univariate trait (see Methods, Supplementary Note).

We collected genome-wide data on open chromatin (inferred from ATAC-seq) and active regulatory regions (inferred from ChIP-seq of histone marks) from a variety of cell-types and tissues, including in-vitro derived CNCCs and their chondrocyte derivatives^49,63^, embryonic craniofacial tissue at different stages of development^64^, neuronal and glial cells from 3D forebrain organoids at various stages of differentiation^50^, and both fetal and adult brain tissue^65^. We first quantified brain and face shape heritability enrichments for these cell-type specific annotations (Supplementary Table 8). Face shape showed significant (5% FDR) heritability enrichment specific to regulatory regions in craniofacial cell-types (mean Z-score 4.58) (Figure 4a). Brain shape showed significant and comparable heritability enrichments for regulatory regions in craniofacial cell-types and tissues, brain organoids, and primary fetal brain tissue (mean Z-scores 4.23, 3.23, 3.33, respectively) (Figure 4b). Within brain organoids, the strongest enrichments were for early-stage glial cells and even earlier-stage whole organoids (mean Z-score 4.11) (Extended Data Fig. 10), consistent with the radial unit hypothesis and in agreement with enrichments of brain surface area heritability^13^. The strong enrichments for craniofacial cell-types, which were substantially more significant than organoid enrichments in the orbitofrontal and medial temporal lobes (Extended Data Fig. 11), suggest that the shared GWAS signal between brain and face shape is mediated primarily by CNCCs and their derivatives early in embryonic development. Consistent with this idea, quantifying brain shape heritability enrichments with the 76 brain-face shared loci removed resulted in decreased enrichment for CNCCs (Z-score difference -0.68) and slightly increased enrichment for the most enriched organoid annotation (Z score difference 0.23) (Extended Data Fig. 12).

**Fig. 4:**
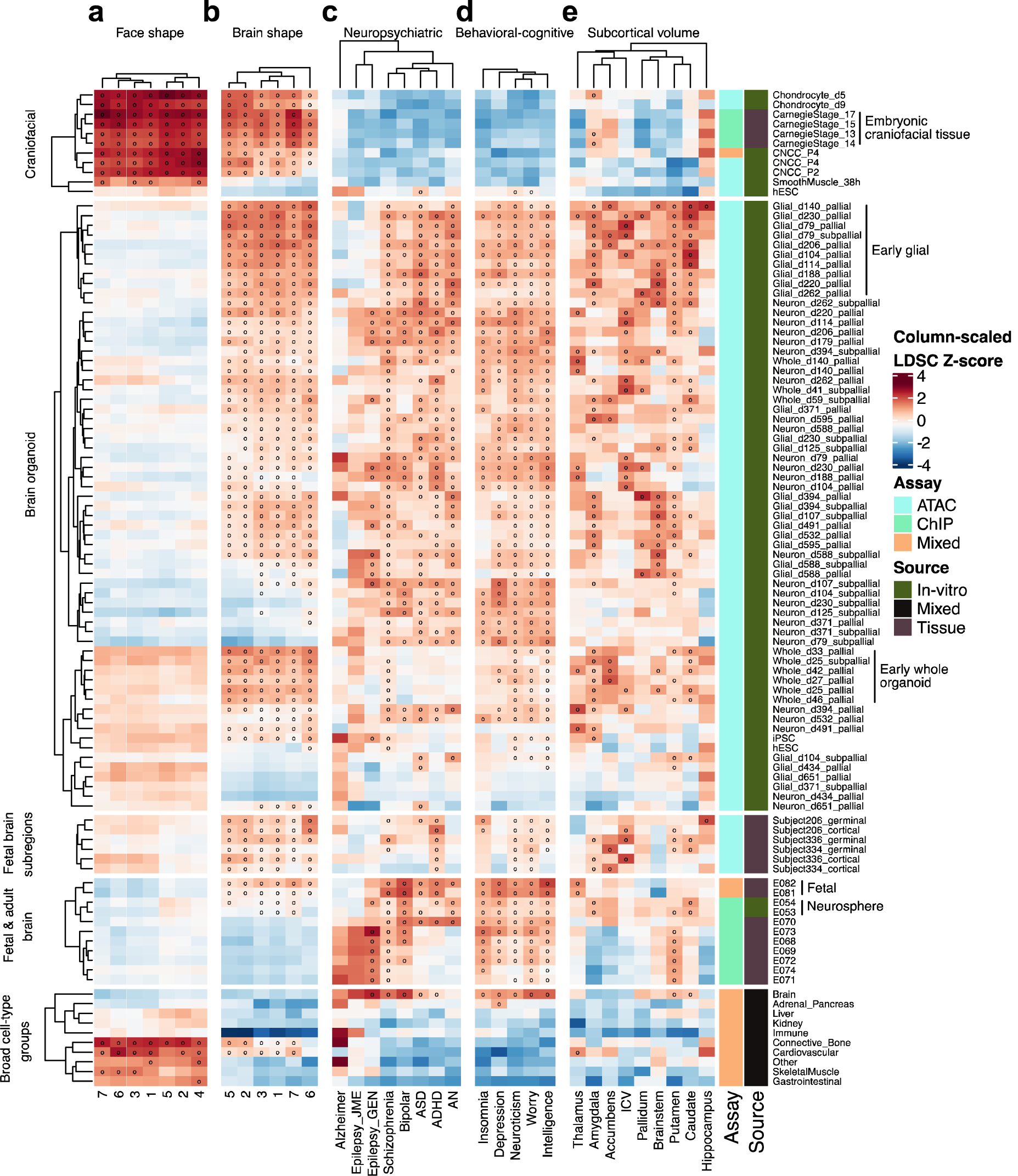
Partitioned heritability enrichments based on cell-type-specific regulatory annotations. Heritability enrichment Z-scores, as estimated by S-LDSC, of **a)** multivariate shape for the first 7 face segments, **b)** multivariate shape for the first 7 brain segments, excluding segment 4 which had low heritability, **c)** neuropsychiatric disorders, **d)** behavioral cognitive traits, and **e)** subcortical volume measures. Heritability enrichments were estimated for annotations based on open chromatin (based on ATAC-seq), regulatory regions (based on ChIP-seq of multiple histone modifications), or a combination of the two. Annotations for the indicated samples, representing in-vitro-derived cell-types, primary tissues, or a combination of both (see Methods for source papers), were added to the S-LDSC baseline model, and the resulting Z-score was scaled by column to visualize relative enrichments between traits. * 5% FDR based on unscaled Z-scores. Trait abbreviations as in Figure 3, with AN representing anorexia nervosa.

Finally, we quantified heritability enrichments for neuropsychiatric disorders, behavioral-cognitive traits, and subcortical volume measures. Neuropsychiatric disorders and behavioral-cognitive traits showed heritability enrichment patterns somewhat distinct from those of brain shape, with significant enrichment for both fetal and adult brain tissue (mean Z-scores 2.17 and 2.64, respectively), and broad enrichment across stages and cell-types of brain organoids (mean Z-score 2.46). In contrast to brain shape, these traits showed no enrichment for craniofacial cell-types or tissues (mean Z-score -0.92) (Figure 4c). Subcortical volume measures showed mixed enrichment patterns, with some regions (amygdala, caudate) similar to those of multivariate brain shape and other regions (putamen) closer to those of neuropsychiatric disorders and behavioral-cognitive traits. These results suggest that while a substantial portion of the shared genetic variation between brain and face shape are mediated by regulatory regions in CNCCs and their craniofacial derivatives, variation in these regulatory regions does not impact neuropsychiatric disorder risk or other behavioral-cognitive traits.

## DISCUSSION

Here, we applied a multivariate phenotyping approach to assess common genetic variation underlying brain shape, revealing a large number of novel loci with effects on brain volume or surface area. While these loci broadly implicate known pathways in brain development, the precise mechanisms by which they modulate brain shape are unknown, suggesting further avenues of investigation. As part of our study, we extended techniques for estimating and partitioning genome-wide heritability, originally developed for univariate traits, to multivariate traits. We anticipate that these and similar extensions will become increasingly useful with the greater availability of high-dimensional imaging or morphological data in large sample sizes.

Our study revealed a striking convergence of common genetic variation affecting brain and face shape, which is at least in part mediated by the regulatory regions active in CNCCs and their more differentiated derivatives. These observations suggest a larger than previously appreciated role of the face in shaping development of the brain and its individual morphological variation. Importantly, however, these shared genetic effects do not appear to significantly impact neuropsychiatric disorder risk or cognitive functions. Our results are therefore consistent with a model whereby CNCCs and their derived cranial structures significantly influence brain shape through both physical interactions and paracrine signaling early in embryogenesis, but later shaping of the cortical morphology, through processes such as gyrification^66^, has a much more significant impact on cognitive traits.

A number of developmental mechanisms could mediate the shared genetics of brain and face shape. One potential contribution comes from the common neuroepithelial origins of the two structures, with genes influencing growth, patterning and cell fate decisions within the neural plate ultimately affecting cell allocation within distinct parts of the brain and face; examples of such neural plate genes within brain-face shared loci include *ZIC2* and *ZIC3*^67–69^. Another potential mechanism entails common genetic variation modulating expression of genes with pleiotropic, independent roles in both brain and face development. A good candidate for this type of mechanism may be the *SOX9* gene, encoding a TF with key functions in neural crest development and chondrogenesis, but which is also required for gliogenesis^70^. Nonetheless, the fact that most brain-face shared genetic effects are concentrated on facial regions from the frontonasal prominence and anterior forebrain regions of the brain suggests additional, proximity-based mechanisms, which can be either structural in nature, or mediated by the paracrine signaling. While development of the brain and face must be tightly coordinated, the brain is thought to have greater structural effects on craniofacial development than the reverse, as the forebrain can act as a structural support for facial development^4^ as well as induce flexion of the basicranium and bone deposition at coronal sutures through tensile forces generated by its growth^3,4,18^. However, we find multiple brain-face shared loci lie near TFs with known, cell-intrinsic roles in, and expression specific to, CNCCs and their derivatives. Furthermore, mutations in these TFs are associated with malformations of the frontal facial skeleton, such as coronal synostosis (*TWIST1*)^45,46^ or fronotonasal dysplasias (*ALX1* and *ALX4*)^43,44^. One possible explanation for these results is that these TFs control regulatory programs that ultimately modulate the ability of the craniofacial skeleton to respond to and accommodate brain growth, thus causing subtle changes in brain shape. It is also possible, however, that these TFs exert some of their phenotypic effects on brain shape by regulating the expression of signaling ligands secreted from the face. For example, CNCCs secrete BMP antagonists which modulate forebrain development by blocking BMP and FGF production in the anterior neural ridge (ANR)^5,6^. BMP antagonist production in CNCCs is regulated by the SIX family TFs^52^, with the *SIX1/4* locus representing one of the shared brain-face signals in our GWAS analyses (Figure 2a). In the reverse direction, studies in chick embryos have shown that Fgf, Shh, and BMP ligands are secreted by the forebrain and regulate the formation of the frontonasal ectodermal zone (FEZ), a signaling center that in turn patterns the frontonasal prominence of the developing face^20–22,71^. Notably, our GWAS analyses implicate all three of these signaling pathways, nominating specific ligands and receptors within those pathways whose modulation may be associated with the brain-face crosstalk. Furthermore, our study nominates other signaling pathways, such as Wnt and TGF-beta, for further investigation in paracrine signaling between the brain and face. Altogether, we uncovered common genetic variants yielding a wealth of candidate molecular players whose diverse mechanistic roles in mediating brain-face interactions during development can be examined in future studies.

Relationships of facial shape with cognitive and personality traits fascinated humans since ancient times, from the Ancient Greeks, who introduced the term ‘physiognomy’ to describe a practice of assessing a person’s personality from their facial appearance^72^, through the Vedic traditions of Samudrika Shastra^73^ and to the Chinese art of face reading^74^. The concept of physiognomy was revived in the late 18^th^ century by Johan Kaspar Lavater, and later gave rise to a related pseudoscientific theory, phrenology, popularized by Franz Josef Gall. Both theories have a troubled history, as they have been used to justify racial discrimination as well as eugenic theories^75,76^. While physiognomy in its original formation has been largely debunked, modern studies have found correlations between facial width-to-height ratios and aggressive tendencies and behaviors^77^, with regrettable renewed efforts in using machine learning approaches to detect such correlations raising serious ethical concerns^78,79^. However, our results argue that while the ancient human intuition of a close relationship between the face and the brain has genetic support at the level of morphology, there is no genetic evidence for the supposed predictive value of face shape in behavioral-cognitive traits, which formed the core of physiognomy and related theories.

## METHODS

### UK Biobank data preprocessing

The UK Biobank project (UKB) is a large dataset of about 500,000 British volunteers with informed consent containing genetics, non-imaging variables and brain imaging data acquired using a fixed protocol^80^. Hereby, brain T1-weighted magnetic resonance imaging (MRI) scans of the UKB, as well as genotyping and covariate information (e.g. sex, age, height, weight, among others), were obtained and used as the discovery dataset. More specifically, we utilized the data release v1.5 of August 2018 which holds a cohort of 21,780 subjects with these three sources of information. This cohort was composed of an adult population (40 to 70 years old, mean of 60 years old), with slightly more females than males (51.6% vs. 48.4% respectively), a predominantly self-reported white British ancestry (97.1%), and an average body mass index (BMI) of 26.6.

For the list of 21,780 subjects, we processed the raw MRI data for a surface-based analysis of the cerebral cortex using the following four-step procedure:

First, the cortical surfaces were segmented and reconstructed from the MRI volumetric data using recon-all (FreeSurfer^81^ *v*.*6*.*0*.*0;* URL section). In this step 20,409 images were processed successfully.

Second, to obtain a minimally preprocessed pipeline similar to the one of the Human Connectome Project (HCP – URL Section) CIFTIFY (Connectivity Informatics Technology Initiative file format - URL section) was used to convert FreeSurfer’s recon-all command output to a HCP-style file format and structure^82^. This protocol converts the data into GIFTI and CIFTI “gray ordinate” file formats, and then performs surface-based alignment of the cortical mesh to the fs_LR Conte69 space using MSMSulc and volume-based registration of subcortical structures to the MNI152 space. High dimensional cortical meshes were down sampled to lower resolution meshes of 32,492 3D vertices (average ∼2mm spacing) and 64,980 triangular faces. Left and right hemispheres were aligned to each other. Due to the alignment of individual brain images with a common brain surface atlas (Conte69), the cerebral cortex was represented by surfaces that were also homologous from one individual to another^83^: a single vertex of a subject’s brain mesh was in very good anatomical correspondence with a single vertex of another subject’s brain mesh, and this for all 32,492 vertices of the meshes. All but one of the images were processed without error in this step.

Third, from the output of CIFTIFY, we selected the mid-cortical surfaces of the left and right hemisphere, which is the surface that runs at the mid-distance between the white surface (which lies at the interface between gray and white matter) and the pial surface (which is the external cortical surface)^84^. The mid-cortical surface does not over or under-represent gyri or sulci^85^, but besides that our choice for this specific surface is arbitrary. The white and/or pial surfaces could have been used alternatively. The vertices from the sub-cortical part of the surface, are typically excluded from surface-based cortex analysis, and were therefore removed based on the sub-cortical vertex index provided by the Conte69 atlas. The final count of vertices for each of the mid-cortical surfaces left and right was 29,759. For each hemisphere and each individual, we also computed the centroid size as the average Euclidean distance of all mesh vertices to the point of gravity, which is a standard measure of size in geometric morphometrics. In the remainder of this work, cortical brain structure or brain shape for short, is represented and referred to by the mid-cortical surface.

Fourth, as quality control for each hemisphere separately, we checked the resulting mid-cortical surfaces for mesh artifacts in a semi-automatic manner. This was done by first measuring the Mahalanobis distance for each individual’s mid-cortical surface to the overall average mid-cortical surface in a generalized Procrustes shape-space spanned by an orthogonal basis of principal components that captures 98% of the total variation. From the distribution of Mahalanobis distances, a z-score for each mid-cortical surface was then established, and each mid-cortical surface with a z-score equal to or larger than three was manually inspected for meshing errors (e.g. triangles stretched too far or triangles folded). All images after step 3 passed this quality control, resulting in a set of 20,407 processed images.

For the list of 20,407 subjects with preprocessed images, we selected genomic data from the UK Biobank, which consisted of the version 3 (March 2018) imputed SNP genotypes, imputed to the Haplotype Reference Consortium and merged UK10K and 1000 Genomes (phase 3) panels. First, European individuals only were selected using principal component analysis (PCA) after excluding SNPs in linkage disequilibrium (LD) from the 1000G Phase 3 dataset (Plink 1.9, 50 variant window-size, 5 variant step size, 0.2 r^2^). A k-nearest neighbor algorithm, using the first 25 reference ancestry principal components, was used to assign a 1000G super population label to each individual, and individuals with the 1000G super population EURO label were selected for analysis only. Second, we filtered the imputed UK Biobank SNPs by removing indels and multi-allelic SNPs, missing genotypes across individuals (<=50%), minor allele frequency (<1%), and Hardy-Weinberg equilibrium (p < 1e^-6^). Third, in order to remove related individuals and capture population structure we pruned the filtered SNP set for LD (Plink 1.9, 50 variant window-size, 5 variant step size, 0.2 r^2^). Subsequently, related individuals were identified and removed when the proportion of identity by descent (IBD) was higher than 0.125. Finally, population structure was captured using principal components analysis (PCA). This ultimately resulted in 9,705,931 filtered SNPs for GWAS analysis on 19,670 unrelated subjects of European descent.

For the list of 19,670 subjects with preprocessed brain and genetic data, we collected the following available list of covariates to control for during statistical testing: genetic sex, age, age-squared, height, weight, diastolic blood pressure, and systolic blood pressure. Additionally, to adjust for population structure the first 20 genetic principal components were included as covariates. Furthermore, the following imaging specific parameters were also included, following Elliot et al.^86^: volumetric scaling from T1 head image to standard space, XYZ-position of brain mask in scanner co-ordinates, Z-position of table/coil in scanner co-ordinates, date of attending assessment center, and assessment center (coded as a dummy variable for each of the 21 centers). For each of the covariate variables, except for assessment center, missing data was replaced by the average value of the respective variables. 26 subjects were removed due to extreme outlying covariate information (>6 times the standard deviation) in weight (11 individuals), diastolic blood pressure (1 individual), systolic blood pressure (3 individuals), X-position of brain mask (6 individuals), Y-position of brain mask (4 individuals), Z-position of table/coil (1 individual). Next, to symmetrize brain shape, the right hemisphere was reflected to the side of the left hemisphere, by simply changing the sign of the x-coordinate for all of the 29,759 3D vertices on the surface of the right hemisphere. Second, we performed a generalized Procrustes superimposition (GPA)^87^, thus eliminating differences in position, orientation, and scale (measured by centroid size) of all left and right hemispheres pooled together. We computed the symmetric brain component as the vertex-wise averaged brain surface of paired and superimposed left and right hemispheres. This resulted in a final discovery dataset of 19,644 subjects containing preprocessed MRI image data on the mid-cortical symmetrized surface, 9,705,931 imputed SNPs and 54 covariates.

### ABCD study data preprocessing

The Adolescent Brain Cognitive Development Study (ABCD) (URL section) is a longitudinal study following brain development and health through adolescence^37^. A total of 11,411 MRI scans with additional information on sex and age were available for download from the data release of April 2019 and of those 11,393 images were processed successfully using the four-step imaging preprocessing described above.

In total 10,627 individuals from the ABCD dataset provided with genetic data on 517,724 SNP variants. These were sent for imputation via the *Odyssey*^88^ pipeline using the SHAPEIT4^89^ and IMPUTE5^90^ workflow to phase and impute respectively. The Haplotype Reference Consortium (HRC)^91^ reference panel was used for imputation. Standard data cleaning and quality assurance practices were performed based on the GRCh37 (hg19) genome assembly. Quality control of the data prior to phasing and imputation includes using the McCarthy Group’s Imputation preparation program (URL section) to check and fix strand, alleles, position, and reference/alternative problems as well as removing ambiguous A/T and G/C SNPS with minor allele frequencies greater than 0.4. Variants that had a missing rate greater than 10% as well as individuals who had more than 10% of missing variants were also excluded from phasing and imputation.

To assess for ancestry, pre-phased quality controlled genotyped variants underwent a filter for Hardy-Weinberg Equilibrium (*P* < 1 x 10^-6^) and were merged with the 1000 Genomes Phase 37 and the Human Genome Diversity Project reference panels. Variants that were in common between the datasets were assessed for LD and then pruned using a 1,500 kb window, 50 bp step size, and a 0.4 r^2^ threshold. This pruned dataset which contained 14,068 individuals from the reference and ABCD datasets were used in a PCA to construct an ancestry space. Using the eigenvalues that were found to explain more than 5% of the total amount of variance, an X-dimensional centroid was created from reference samples designated as having European ancestry. This in term created a “European centroid.” Only participants that were within 3 standard deviations of the centroid were retained to obtain a relatively homogenous sample. Following QC and ancestry assessment the ABCD dataset was trimmed down to 5,622 individuals and 484,000 variants. Following phasing and imputation, variants were filtered based on the imputation quality control INFO metric (INFO score > 0.7), which resulted in 15.3M imputed ABCD variants.

For the 5,622 individuals of primarily European ancestries, the genotyped and imputed variants were filtered by removing indels and multi-allelic SNPs, missing genotypes across individuals (<50%), minor allele frequency (<1%), and Hardy-Weinberg equilibrium (p < 1e^-6^). Subsequently, in order to remove related individuals and capture population structure we pruned the filtered genotyped SNP set for LD (Plink 1.9, 50 variant window-size, 5 variant step size, 0.2 r^2^). Subsequently, 1,009 related individuals were identified and removed when the proportion of identity by descent (IBD) was higher than 0.125. Finally, population structure was captured using PCA. Of the 4,613 unrelated subjects of European ancestries, 143 did not have a preprocessed brain image. This resulted in a final replication dataset of 4,470 individuals with preprocessed MRI image data, representing brain shape, 15.3M imputed SNPs and 7 covariates (sex, age and the first 5 genetic PCs). The minimum and maximum age of this final replication dataset, was 8.9 years and 11 years, respectively, with a mean age of 9.9 years. 46.5% are female and 53.5% are male.

### Auxiliary traits GWAS summary statistics

We collected publicly available genome-wide summary statistics for 22 auxiliary traits encompassing neuropsychiatric disorders^92–97^, behavioral-cognitive traits^98–100^, and subcortical volume measures^33–35^. In Supplementary Table 7, we provide links to relevant publications and URLs for these summary statistics.

### Point-wise SNP-heritability estimation of the mid-cortical surface

For each of the 29,759 vertices of the averaged mid-cortical 3D surfaces in the UK Biobank we computed a multivariate (X, Y and Z coordinate per vertex) narrow-sense heritability from common SNP variants using a linear mixed model (LMM). A genomic relationship matrix (GRM) modelled as random effects in the LMM was computed from LD pruned SNP data (Plink 1.9, 50 variant window-size, 5 variant step size, 0.2 r^2^). The first 10 genomic principal components and additional covariates (sex, age, height, weight, diastolic and systolic blood pressure) were modelled as fixed effects in the LMM. We used the open-source software SNPLib (URL Section)^101^, whose implementation is equivalent to the widely used GCTA software^102^ for a homogenous population.

### Global-to-local (G2L) segmentation of the mid-cortical surface

The UK Biobank (n=19,644) served as discovery cohort using a data-driven global-to-local (G2L) segmentation of brain shape similar to previous work on face shape^15,16^. First, the superimposed and symmetrized mid-cortical surfaces were corrected using a partial least-squares regression (PLSR, function plsregress from Matlab 2019b) for all UK Biobank covariates listed above, augmented with centroid size to eliminate allometric effects of size on brain shape^87^. Second, pair-wise structural connections based on the multivariate generalization of the Pearson correlation, or RV-coefficient^103^, between each pair of 3D surface vertices generated a squared similarity matrix (29,759 x 29,759). Third, a Laplacian transformation was applied to enhance similarities prior to an eigendecomposition of this squared matrix. Finally, within the eigen spectral map, K-means++ clustering was used to group highly correlated vertices, that, when mapped back to the brain surface, result in a segmentation of the brain into separate modules. This was done in a bifurcating hierarchical manner using eight levels, resulting in a total of 511 hierarchically linked facial segments, with 1, 2, 4, 8, 16, 32, 64, 128, 256 non-overlapping modules at levels 0, 1, 2, 3, 4, 5, 6, 7, 8. In contrast to our work on facial shape^15,16^, we pruned down segments with fewer vertices than 1% of the total vertex count. I.e., segments with less than 30 vertices were removed as to safeguard the minimum size of each segment. This resulted into the pruning of 226 segments, generating a final G2L segmentation of brain shape consisting of 285 segments across eight levels as depicted in Figure 1. The hierarchical design provided a shape decomposition focused at different levels of detail, going from the full hemisphere, and larger brain segments, to more local smaller brain segments. This allowed the investigation of localized shape variations on the one hand, towards larger, more integrated shape variations on the other hand.

For each of the 285 brain segments separately, the group of surface vertices in a segment were subjected to a new GPA. As such, a multivariate shape-space for each brain segment was constructed independently of the other segments and its relative positioning within the full hemisphere. Subsequently, after GPA, each segment’s shape-space was spanned by a multivariate orthogonal basis using PCA on the pooled x, y and z coordinates of the collection of superimposed vertices in that segment. Finally, we retained enough PCs to explain up to 80% of the total shape variation within each segment. This is in slight contrast to our previous work on facial shape, where we used a parallel analysis (PA) instead. By choosing those PCs explaining up to 80% of the variation we typically retained 50% of the components otherwise retained using PA (e.g. 437 instead of ∼1000 for the full hemisphere). However, the number of components retained using PA became computationally intractable in a GWAS context. Therefore, we opted to further reduce the number of PCs per brain segment, knowing that these certainly represent non-noisy shape variations, confirmed by the PA.

### Overlap of brain atlases with G2L segmentation

We investigated the overlap of brain segments at each of the eight levels from our G2L segmentation with brain regions from three commonly used brain atlases (Desikan Killiany (34 distinct gyral based regions)^29^, Destrieux (74 distinct gyral and sulcal based regions)^30^, and the Glasser (180 distinct multi-modal based regions)^31^) that were also defined within the HCP project data format and thus defined on the same surface mesh of 29,759 vertices. For each of the G2L levels separately, every brain surface vertex has a unique label of the G2L brain segment it belongs to at that level and a unique label of the atlas brain region it belongs to. Using these two labels per vertex the normalized mutual information across all vertices provided a measure of overlap from 0 (no overlap) to 1 (complete overlap), for each G2L level with each of the three atlases. Additionally, each brain segment and each brain region defined a subset of vertices, and therefore, for each segment we defined the intersection of vertices with each brain region, and for each brain region we defined the intersection of vertices with each brain segment, expressed as percentages.

### G2L multivariate genome-wide discovery

The global-to-local phenotyping partitioned cortical brain shape into overlapping (across different hierarchical levels), as well as non-overlapping (within a single hierarchical level) segments, each of which was represented by a different subset of mid-cortical surface vertices and spanned by multiple dimensions of variation (principal components, PCs). For each brain segment separately, canonical correlation analysis (CCA, canoncorr from Matlab 2019b), was therefore used as a multivariate testing framework (note that CCA is also implemented in Plink 1.9 for multivariate phenotypes). CCA extracts the linear combination of PCs spanning the brain segment that correlates maximally with the SNP variant being tested, and therefore reveals a latent shape trait within the shape-space of the brain segment. The correlation of this latent shape trait with the SNP variant is tested for significance based on a Chi-squared (X^2^) statistic (right-tail, one-sided test), with degrees of freedom equal to the dimensionality or number of principal components of the brain segment under investigation. Using CCA, we tested each SNP (n=9,705,931) individually under an additive genetic model in the UK Biobank (n=19,670) against each of the brain segments (n=285) separately. Note that CCA does not accommodate adjustments for covariates, but effects of important covariates were corrected for (using PLSR) at the phenotyping stage. Additionally, we applied a similar correction for the covariates on each SNP, again using PLSR, excluding this time the covariates that were only relevant for the correction of imaging data (e.g. acquisition center). Therefore, the CCA analysis was performed under the reduced model, which was obtained after removing the effects of covariates on both the independent SNP variants as well as the dependent multivariate brain shape phenotypes.

Given the burden of multiple comparisons, a strict significance threshold of *P* ≤ 5 x 10^-8^ was used to declare “genome-wide significance”, which corresponds to a Bonferroni correction for 1 million independent tests and is mostly applicable in a GWAS on a European-ancestry cohort^104^. Due to 285 multivariate GWAS runs, the multiple comparisons burden was magnified. Therefore, we also determined a more stringent threshold for declaring “study-wide significance” corresponding to an additional adjustment for the effective number of independent tests. In a first instance, looking at the number of eigenvalues larger than one of a pairwise multivariate correlation (RV-coefficient) matrix (285 x 285) across all segments^105^, determined a total of 210 independent tests. In a second instance, an empirical estimate of the number of independent tests was also obtained using the 472 lead SNPs representing the genome-wide associated independent genetic loci (see description below), to keep the estimations computationally tractable. First, for a single SNP we randomly permuted the genotypes in the UK Biobank, essentially creating genotypes that have a noisy association with multivariate brain shape. Then, we performed the CCA associations of the randomized genotypes to each of the 285 brain segments and retained the lowest or “best” p-value out of the 285 p-values obtained. Step 1 and 2, were repeated 10,000 times. Subsequently, we divided 0.05 by the 5th percentile of the 10,000 permuted best CCA p-values, and this was done for each of the 472 SNPs. Based on these 472 outcomes, the mean estimation of the number of empirical independent tests based on 285 brain segments is 241.46 (11.09 standard deviation). Because the empirical estimation was more conservative compared to the eigenvalue-based estimation we opted for the former and determined the study-wide significance threshold to be 2.0707 ⨯ 10^-10^ (i.e., 5 ⨯ 10^-8^ / 241.46).

### Peak detection, overlap and annotations

We observed 38,630 SNPs and 23,413 SNPs at the level of genome-wide and study-wide significance, respectively. These were clumped into 472 (genome-wide) and 243 (study-wide) independent loci in three steps. First, starting with the best associated or lead SNP (lowest p-value), SNPs within 10kb or within 1Mb but with r^2^ > 0.01 were clumped into the same locus represented by the lead SNP. This process was repeated until all SNPs were assigned into 509 loci. Second, based on the lead SNPs only, a wider window of +/- 10Mb was tested for r^2^ > 0.01, reducing the number of loci (n=502) by merging seven lead SNPs. Third, any locus with a singleton lead SNP (without additionally clumped SNPs) below the study-wide threshold was removed (n=30). r^2^ values were computed using the genotypes from the UK Biobank.

To study the functional enrichment for genes near the 472 genome-wide lead SNPs, we performed gene ontology (GO) analysis using GREAT^40^ (v4.0.4) and FUMA^39^ (v1.3.6) using default settings. GO terms that were significant by both binomial and hypergeometric tests (False Discovery Rate (FDR) q-value < 0.05) across three or two windows were reported as strongly and weakly enriched respectively.

In determining overlap between lead SNPs from different GWAS, we used a similar strategy: two lead SNPs tag the same genetic locus if they are within 10kb of each other or if they are within 1Mb of each other and with r^2^ > 0.2. For considering and quantifying overlap between the 472 brain shape loci and 430 other studies from the NCBI-EBI GWAS Catalog, we defined LD blocks of 0.2 around the 472 loci using Plink 1.9, and then calculated odds ratio and *P* for the overlap between these blocks and any given GWAS using bedtools v2.27.1, fisher function.

In determining brain-face shared loci, we first started from the 472 genome-wide lead SNPs from the brain GWAS and looked for any SNP within 10kb or within 1Mb and LD > 0.2 of these lead SNPs with at least a genome-wide suggestive (*P* ≤ 5 ⨯ 10^-7^) association in the facial multivariate GWAS^16^. This resulted in 57 loci with evidence of association in brain and face shape. Then we took the 203 genome-wide lead SNPs reported in the face GWAS^16^, and clumped them if two lead SNPs were within 10kb or within 10Mb but with r^2^ > 0.01. For the resulting 197 independent genome-wide facial lead SNPs we selected any SNP within 10kb or within 1Mb and with r^2^ > 0.2 with at least a genome-wide suggestive (*P* ≤ 5 ⨯ 10^-7^) association in the brain shape GWAS. This resulted in another 54 loci with evidence of association in brain and face shape and together with the previous 57 loci they were clumped (within 10kb or within 1Mb and r^2^ > 0.2) into a final set of 76 independent brain-face shared loci.

We identified candidate genes in the vicinity of the 76 brain-face shared loci through a manual process. For each locus, we first considered all genes within 500kb of the lead SNP. We primarily relied on evidence for these genes’ involvement in a human craniofacial or neurodevelopmental syndrome, or for evidence of craniofacial or neurodevelopmental defects in knockouts of their orthologs in mice. Secondarily, we also considered associations with Gene Ontology (GO) terms related to craniofacial development, neurodevelopment, or skeletal system development. In some cases (i.e. *SOX9*, where enhancer-promoter interactions over 1Mb have been described^49^), we extended the window to within 750kb of the lead SNP.

### ABCD replication testing

The ABCD study data (n=4,470) was used as replication panel, with the UK Biobank discovery cohort used as a phenotyping reference. First, the GPA superimposed and symmetrized mid-cortical shapes were corrected for the confounders of sex, age, and the first five genomic PCs, augmented with centroid size to eliminate allometric effects of size on brain shape^87^ using PLSR. Second, the PLSR residuals that were centered on the overall average brain shape of the ABCD study, were added to the overall average brain shape of the UK Biobank. Third, the corrected and re-centered brain shapes were segmented using the G2L segmentation and projected onto the principal components of the segments from the UK Biobank. This ensured consistency in brain segment delineation and shape-space across both datasets.

For a particular discovery lead SNP in a particular brain segment the replication panel was projected onto the latent shape trait of the lead SNP. This generated univariate projection scores as phenotypes^106^ to test for in the replication panel that are equivalent to the latent shape traits or phenotypes in the discovery panel, i.e. the latent shape trait, once discovered using CCA, was fixed and explicitly measured in the replication cohort. Replication was therefore tested using a standard univariate linear regression (two-sided, regstats Matlab 2019b). This was done for each of the 466 lead SNPs for which the exact SNP or a proxy SNP (within 10kb or within 1Mb and r^2^ > 0.2) was available for analysis in the ABCD cohort, and in each of the 285 segments that were associated at *P* ≤ 5 x 10-8, which resulted in 3,586 replication tests. From all replication efforts combined (n=3,586), we computed a 5% FDR-adjusted significance threshold^107^ equal to *P* ≤ 0.0369.

### Clinical gene-panel overlap

Gene panels were downloaded from the Genomics England PanelApp website. Only panels that will be used for clinical interpretation in the 100,000 Genomes project were selected (provided by PanelApp^41^). The clinical gene-panels were merged in disease (sub)categories according to the 100,000 Genomes Project criteria (e.g. the clinical gene panel “Intellectual Disability” belongs to the sub-category “Neurodevelopmental Disorders”, which is part of the “Neurology and Neurodevelopment” disease category). Only genes with a high level of confidence for gene-disease association were included in the clinical gene panels. We calculated the overlap between genes from clinical panels/subcategories/categories and different gene-sets allowing for a 200kb, 500kb or 1Mb window around the loci. Permutation testing was done to see if this overlap was higher than expected by chance. In brief, we generated 10,000 random panels for each clinical panel/subcategory/category with equal size using a list of 19,198 protein-coding genes. P-values were obtained by dividing “the number of times the overlap random panel and gene-set was larger than the overlap clinical gene-panel/subcategory/category and gene-set” and “number of random gene-panels created (10,000)” . Clinical panels/subcategories/categories were interpreted as strongly or weakly enriched if they showed significance (*P* < 0.05) across three or two different gene-sets respectively.

### Expression analyses of candidate genes at brain-face overlapping loci

Gene expression levels (log2(TPM) values) for three-dimensional forebrain organoids and purifying neuronal or glial lineages were obtained from Trevino et al^50^ (GSE132403). Raw RNA-seq reads from CNCCs at passages 1-4, as well as day 9 chondrocytes derived from P4 CNCCs, were obtained from Long et al^49^ (GSE145327), and TPM values were quantified using kallisto^108^ with sequence-biased bias correction enabled.

### LD score regression SNP-heritability for multivariate traits

In the Supplementary Note, we provide a general proof showing that when applying LD score regression (LDSC) to summary statistics of a multivariate GWAS, albeit with a small correction to the resulting χ^2^ statistics, the heritability estimated by the LDSC slope is equal to 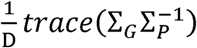, which is a *D*-dimensional generalization of heritability for genetic and phenotypic covariance matrices Σ_G_ and 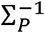, respectively. When the dimensions of the multivariate trait are either genetically or phenotypically uncorrelated, as is the case in both the brain and face GWAS, this expression simplifies to the average SNP-heritability across dimensions, but we also demonstrate extensions for correlated dimensions. Similarly, when applying stratified LD Score regression (S-LDSC), one obtains enrichments for this partitioned average heritability. We further show that 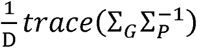 is an appropriate multivariate generalization of heritability since it is the only measure to satisfy the following four properties: 1) invariance to units of measurement, 2) coordinate-free, 3) linear in Σ_G_, and 4) maximized with a value of 1 when 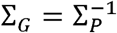.

Thus, for brain and face shape, we applied LDSC and S-LDSC using the published software (URL section) to corrected χ^2^ statistics from GWAS of each brain or face segment, since a full multivariate GWAS was performed in each segment. We used unmodified chi-squared values for the univariate traits analyzed (including indicated cases where we performed individual, univariate GWAS for each PC in brain and face segment 1). While using unmodified chi-square values results in a small bias, we used unmodified statistics for univariate traits for consistency with previous studies. For S-LDSC analyses, we limited ourselves to traits with SNP-heritability Z-scores > 7, as done in Finucaine et al^62^, unless otherwise indicated.

### Functional annotations for stratified LD score regression

We downloaded a range of publicly available cell-type and sample-specific annotations representing open chromatin and/or active regulatory regions. Specifically, we obtained data on open chromatin (all ATAC-seq peaks) from brain organoids^50^, fetal brain tissue^109^, and CNCCs and derived chondrocytes^49^. Raw ATAC-seq reads from Long et al were mapped to hg19 with bowtie2^110^ with default settings, and peaks were called using MACS2^111^ with default settings. Annotations for active regulatory regions (based on a range of epigenomic marks) were obtained from CNCCs^63^, embryonic craniofacial tissues^64^, fetal and adult brain tissue^65^, and broad groupings of cell-types^62^. For CNCC data from Prescott et al^112^, we combined all regions annotated as enhancers (weak, intermediate, strong) or promoters (weak and strong); For embryonic craniofacial tissues, we combined all regions with the following annotations from the 25-state chromHMM model: ‘Enh,’ ‘TxReg,’ ‘PromD1,’ ‘PromD2,’ ‘PromU,’ ‘TssA.’ For fetal and adult brain tissue, we combined all regions with the following annotations from the 15-state chromHMM model: ‘1_TssA,’ ‘2_TssAFlnk,’ ‘7_Enh,’ ‘6_EnhG.’ Each annotation was individually added to the baselineLD model from Finucaine et al, which includes various annotations of genes, conserved regions, and general enhancers and promoters. The resulting S-LDSC output (heritability fold-enrichment magnitude and significance, as well as coefficient Z-scores) are provided in Supplementary Table 8. When quantifying heritability enrichments with brain-face shared loci removed, we removed all SNPs within the same approximately independent LD block^113^ as one of the 76 brain-face shared loci and re-computed LD scores as well.

### Quantifying sharing of signals between pairs of GWAS

To assess the extent to which genome-wide profiles of association were shared between a pair of phenotypes (e.g. multivariate brain and/or face segments and/or univariate neuropsychiatric, behavioral-cognitive, and subcortical volumes), we computed a Spearman correlation between two vectors of LD-block organized association p-values. First, as is done in LDSC, genome-wide SNPs were selected to overlap with the HapMap3 SNPs^114^ and SNPs within the major histocompatibility complex (MHC) region were removed. Second, we downloaded the locations of 1,725 blocks in the human genome that can be treated as approximately independent LD blocks in individuals of European ancestries^113^, and organized all SNPs within these blocks. For every LD block we computed the mean SNP –log10(p-value), and then computed a rank-based Spearman correlation using the averaged association values (n=1,725) for each LD block. A standard error of the Spearman correlation was estimated using statistical resampling with 100 bootstrap cycles with replacement from the 1,725 LD blocks.

## Supporting information

SI Guide

## Ethics statement

This study was conducted in compliance with the principles of the Declaration of Helsinki, the principles of GCP and in accordance with all applicable regulatory requirements. Local ethics review and approval for this study (S63179) was performed and obtained from the ethical committee for research of the University Hospital UZ Leuven and the University KU Leuven.

## URLs

UK biobank: https://www.ukbiobank.ac.uk/about-biobank-uk/

Human Connectome Project: http://www.humanconnectomeproject.org/

Adolescent Brain Cognitive Development (ABCD) study: https://abcdstudy.org/about/

SNPLIB: https://github.com/jiarui-li/SNPLIB

Freesurfer: https://surfer.nmr.mgh.harvard.edu/

Ciftify: https://github.com/edickie/ciftify and https://www.nitrc.org/projects/cifti/

Conte69 Atlas: http://brainvis.wustl.edu/wiki/index.php//Caret:Atlases/Conte69_Atlas

McCarthy Tools. https://www.well.ox.ac.uk/~wrayner/tools/

LDSC: https://github.com/bulik/ldsc/wiki

## Data availability

All the data and detailed information for the UK Biobank, including genetic markers, covariates and MRI images are available to bona fide researchers via the UK Biobank data access process (see http://www.ukbiobank.ac.uk/register-apply/).

All the data and detailed information for the ABCD study, including genetic markers, covariates and MRI images are also available to bona fide researchers through the ABCD data depository (https://nda.nih.gov/abcd/request-access)

Relevant data and materials from the facial GWAS study are available online (https://doi.org/10.6084/m9.figshare.c.4667261). The full facial GWAS summary statistics will be uploaded to GWAS catalog (https://www.ebi.ac.uk/gwas/) upon publication of that work that is currently formerly accepted at Nature Genetics. The facial GWAS paper is currently accessible on bioRXiv (https://www.biorxiv.org/content/10.1101/2020.05.12.090555v1). Furthermore, relevant files generated from the face and brain GWAS summary statistics as input to (S-)LDSC regression and spearman correlations are available on FigShare, see table below. The full brain GWAS summary statistics will be uploaded to the GWAS Catalog upon publication.

All relevant additional data related to this work are provided in the FigShare repository for this work (https://figshare.com/s/bfc96fe9375e9f787b4b). This includes additional figures, input files and updated implementations, listed in the table below.

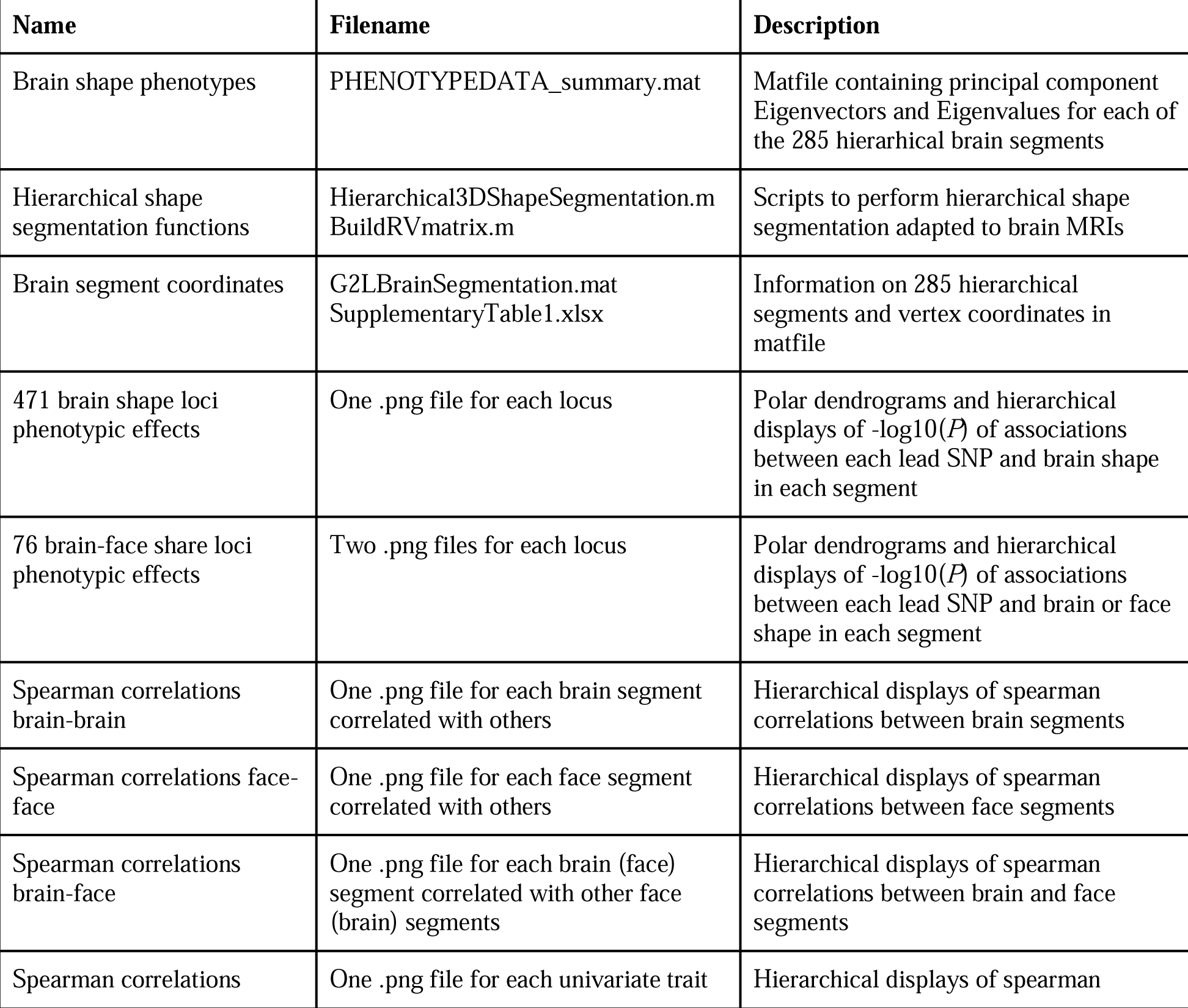
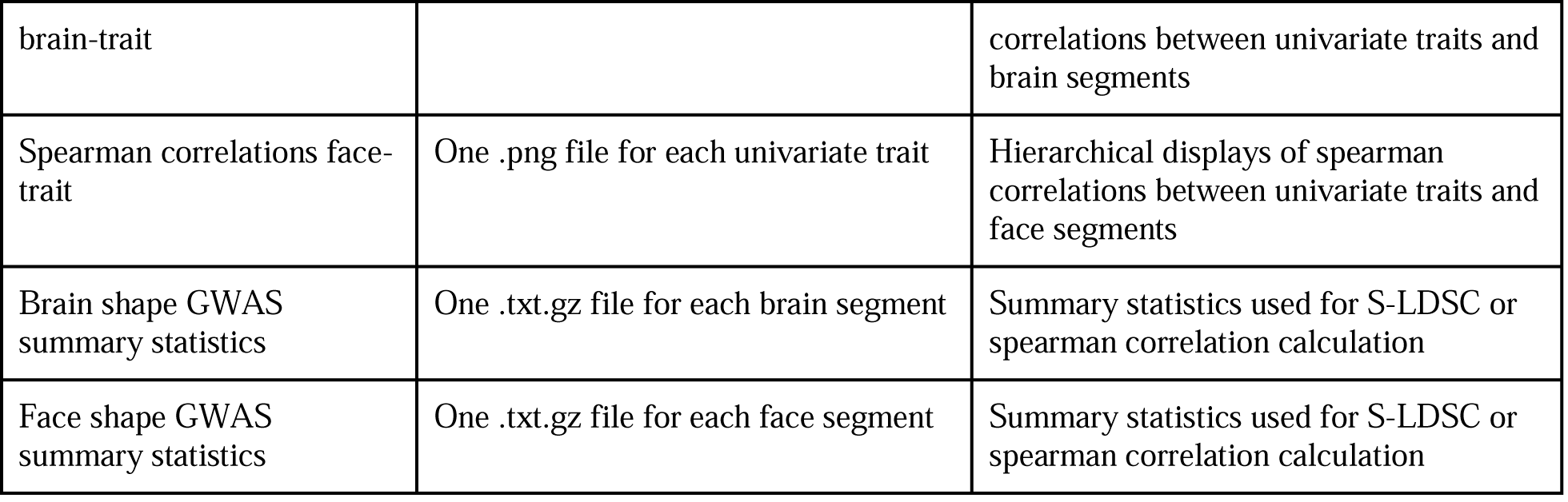

## Code availability

Matlab implementations of the hierarchical spectral clustering to obtain phenotypic shape segmentations are available from a previous publication https://doi.org/10.6084/m9.figshare.7649024.v1). Updated implementations used in this work are provided as described in the Figshare table above. The statistical analyses in this work were based on functions of the statistical toolbox in Matlab as mentioned throughout the Methods. Other materials and external software used mentioned throughout the Methods are available online (see URL section).

## Acknowledgements

JW was supported by the Howard Hughes Medical Institute, a Lorry Lokey endowed professorship and a Stinehart Reed award, SN was supported by a Helen Hay Whitney Fellowship. The KU Leuven research team and analyses were supported by the National Institutes of Health (1-R01-DE027023, 2-R01-DE027023), The Research Fund KU Leuven (BOF-C1, C14/15/081 & C14/20/081) and The Research Program of the Research Foundation - Flanders (Belgium) (FWO, G078518N). The computational resources and services used in this work were provided by the VSC (Flemish Supercomputer Center), funded by the Research Foundation - Flanders (FWO) and the Flemish Government – department EWI. JPS was supported by a National Institutes of Health training grant (5T32HG000044-23). JKP was supported by the National Institutes of Health (HG008140, HG009431). Pittsburgh personnel, data collection, and analyses were supported by the National Institute of Dental and Craniofacial Research (U01-DE020078, R01-DE016148, and R01-DE027023). Funding for genotyping by the National Human Genome Research Institute (X01-HG007821 and X01-HG007485) and funding for initial genomic data cleaning by the University of Washington provided by contract HHSN268201200008I from the National Institute for Dental and Craniofacial Research awarded to the Center for Inherited Disease Research (https://www.cidr.jhmi.edu/). JCT was supported by the National Institutes of Health (5R01-DA033431-07) and the National Science Foundation (1922598).

This research has been conducted [in part] using the UK Biobank Resource under application number 43193 (understanding the genetic architecture of human brain shape from MRI using global-to-local shape segmentations), and we are grateful for all the participants in that resource. This manuscript reflects the views of the authors and may not reflect the opinions or views of the UK Biobank funders and investigators.

Data used in the preparation of this article were obtained from the Adolescent Brain Cognitive Development (ABCD) Study (https://abcdstudy.org), held in the NIMH Data Archive (NDA). This is a multisite, longitudinal study designed to recruit more than 10,000 children age 9-10 and follow them over 10 years into early adulthood. The ABCD Study is supported by the National Institutes of Health and additional federal partners under award numbers U01DA041022, U01DA041028, U01DA041048, U01DA041089, U01DA041106, U01DA041117, U01DA041120, U01DA041134, U01DA041148, U01DA041156, U01DA041174, U24DA041123, U24DA041147, U01DA041093, and U01DA041025. A full list of supporters is available at https://abcdstudy.org/federal-partners.html. A listing of participating sites and a complete listing of the study investigators can be found at https://abcdstudy.org/Consortium_Members.pdf. ABCD consortium investigators provided data but did not necessarily participate in analysis or writing of this report. This manuscript reflects the views of the authors and may not reflect the opinions or views of the NIH or ABCD consortium investigators.

---

**Extended Data Fig. 1.**
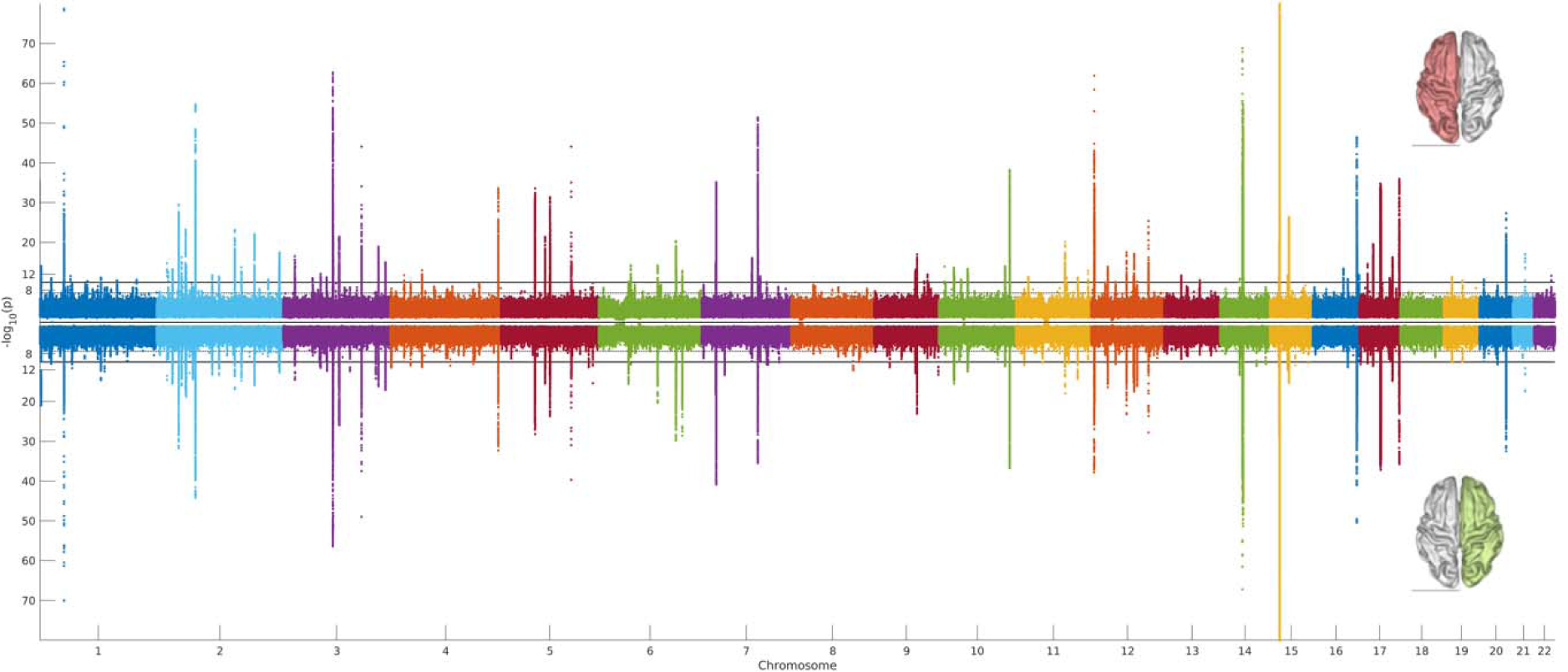
Miami plot of brain shape in left (top) and right (bottom) hemispheres in the UK Biobank. Global-to-local segmentation and CCA-based GWAS was performed independently within each hemisphere. For each SNP, the aggregated p-values across all left (n=241) or right (n=201) hemisphere segments are plotted.

**Extended Data Fig. 2.**
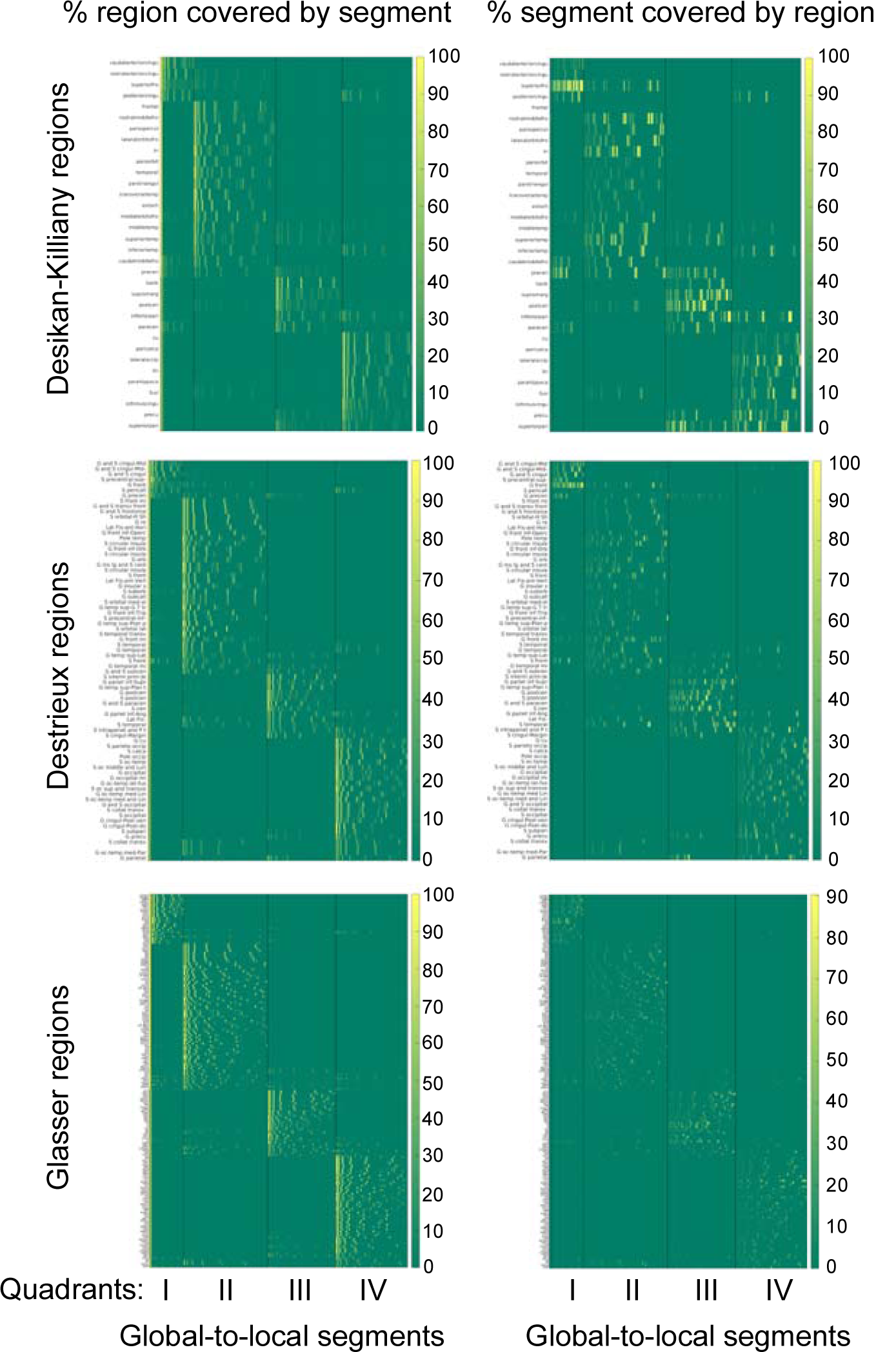
Overlap of global-to-local segmentation of brain shape with commonly used brain atlases.

**Extended Data Fig. 3.**
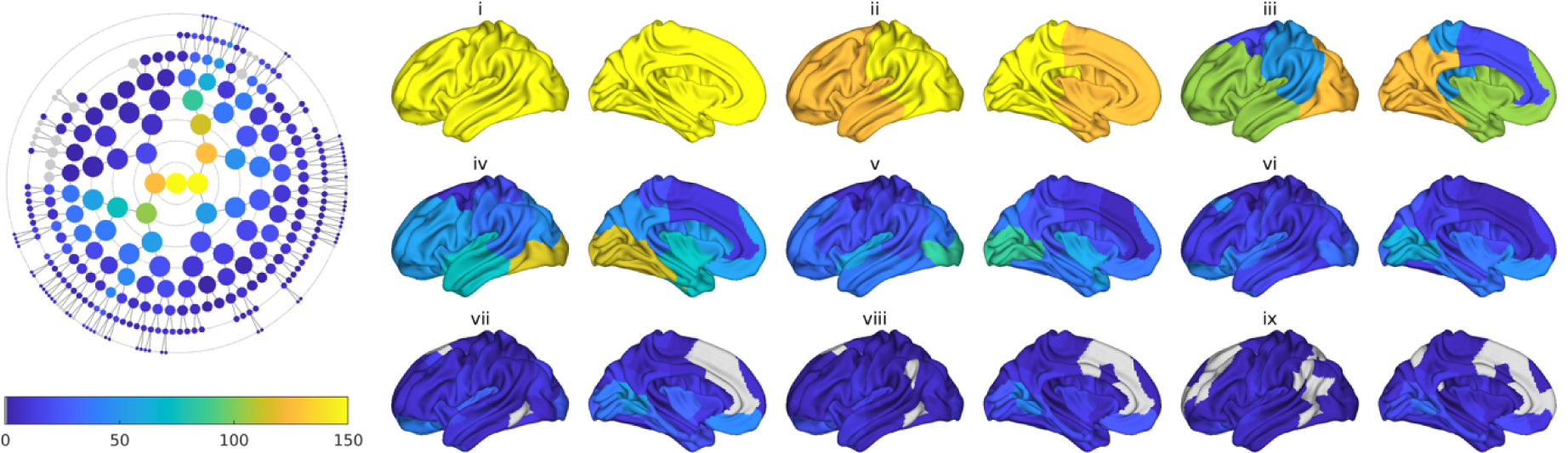
Number of independent genome-wide significant associations discovered by hierarchical segments. Among segments of each hierarchical level (indicated by lower-case Roman numerals and corresponding to concentric circles in the polar dendrogram), the number of independent genome-wide significant associations is shown.

**Extended Data Fig. 4.**
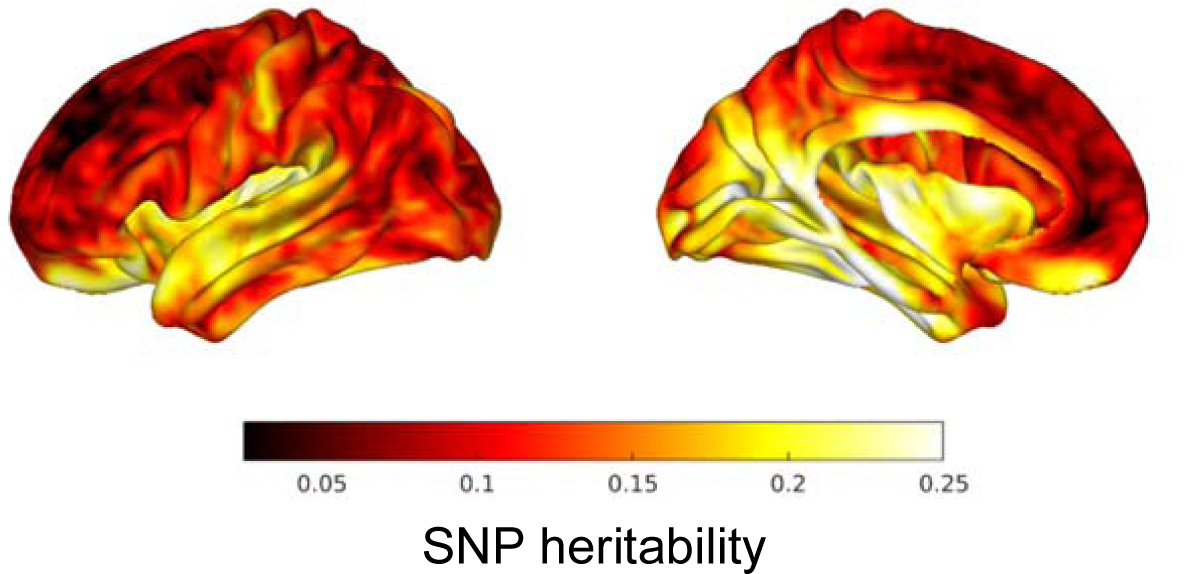
Point-wise SNP heritability estimates across the mid-cortical surface.

**Extended Data Table 1.** Overlap between previous GWAS of brain surface areas or subcortical volumes with brain shape GWAS in this study. ‘Subcortical combined’ refers to a combined set of loci from four studies of subcortical volume measures^32–35^.

| Study | # loci tested | # lead SNP $P < 5 \times 10^{-8}$ | # proxy SNP $P < 5 \times 10^{-8}$ | % overlap |
| --- | --- | --- | --- | --- |
| Subcortical combined<sup>31-34</sup> | 65 | 15 | 18 | 27.6 |
| Grasby et al<sup>13</sup> | 301 | 195 | 236 | 78.4 |
| Zhao et al<sup>35</sup> | 494 | 212 | 273 | 55 |

**Extended Data Fig. 5.**
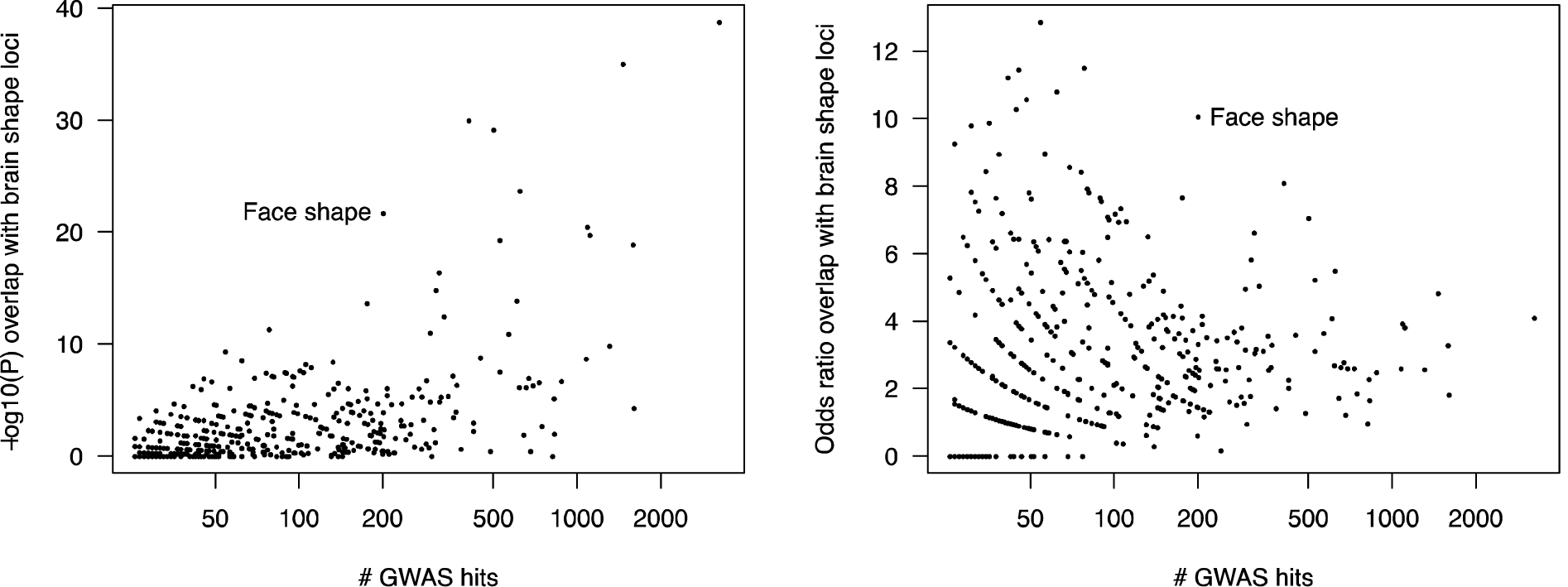
Overlap between genome-wide significant brain shape loci and genome-wide significant loci from 430 other studies. GWAS hits (number on x-axis) for other studies were obtained from the NCBI-EBI GWAS Catalog, and *P*-values (left, y-axis) and odds ratios (right, y-axis) for significance of overlap with regions in LD (> 0.2) with brain shape loci were computed using bedtools’ fisher function (see Methods). Note that relative to other traits with equivalent numbers of GWAS hits, face shape shows overlap with brain shape loci greater in both significance and magnitude than.

**Extended Data Fig. 6.**
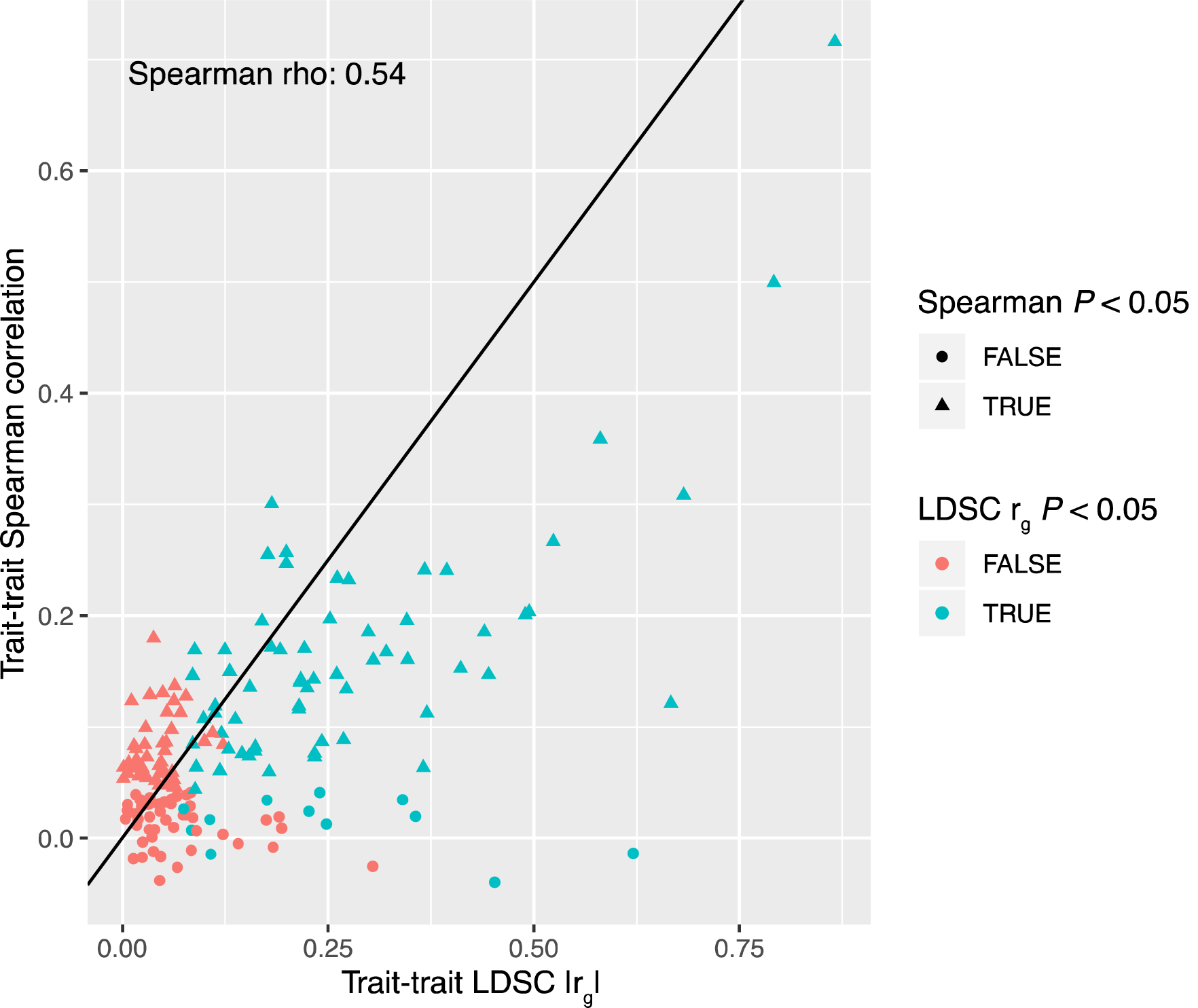
Comparison of LDSC genetic correlations and Spearman correlation between pairs of univariate traits. Each point represents a pair of univariate traits (of all those considered in this study, see Methods), while the x- and y-axes indicate the absolute value of the LDSC-estimated genetic correlation and the estimated genome-wide sharing of effects by the Spearman correlation method. Point colors and shapes indicate significance (*P* < 0.05) from LDSC or the Spearman correlation method, respectively.

**Extended Data Fig. 7.**
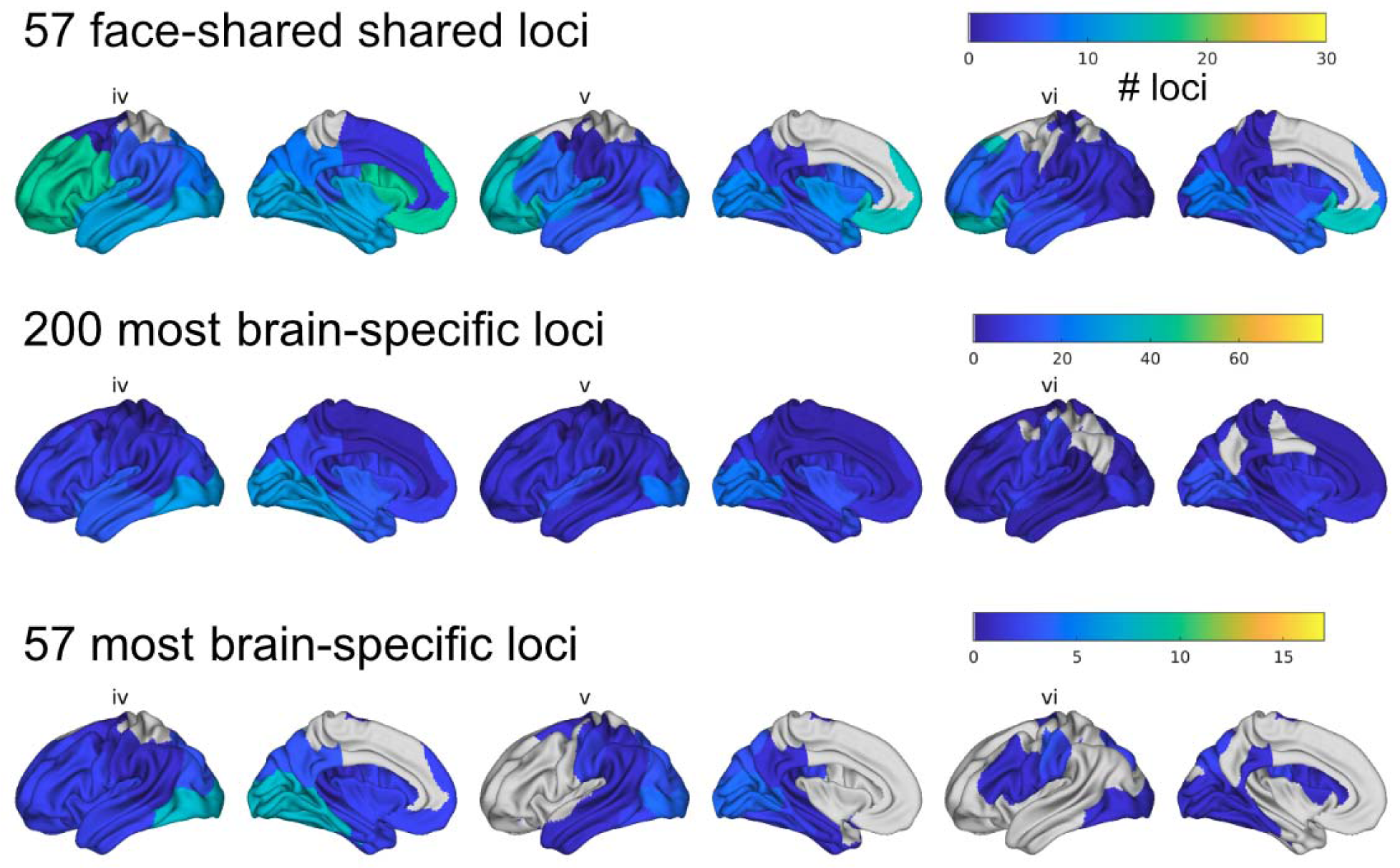
Regional associations of genome-wide significant loci for brain shape stratified by shared effects on facial shape. For the indicated sets of genome-wide significant brain shape loci, the number of associations with each brain segment (shown at hierarchical levels iv-vi) was plotted. The top and bottom 57 face-shared or brain-specific loci were chosen as 57 is the number of brain shape loci which have at least suggestive (*P* < 5×10^-7^) association with face shape.

**Extended Data Fig. 8.**
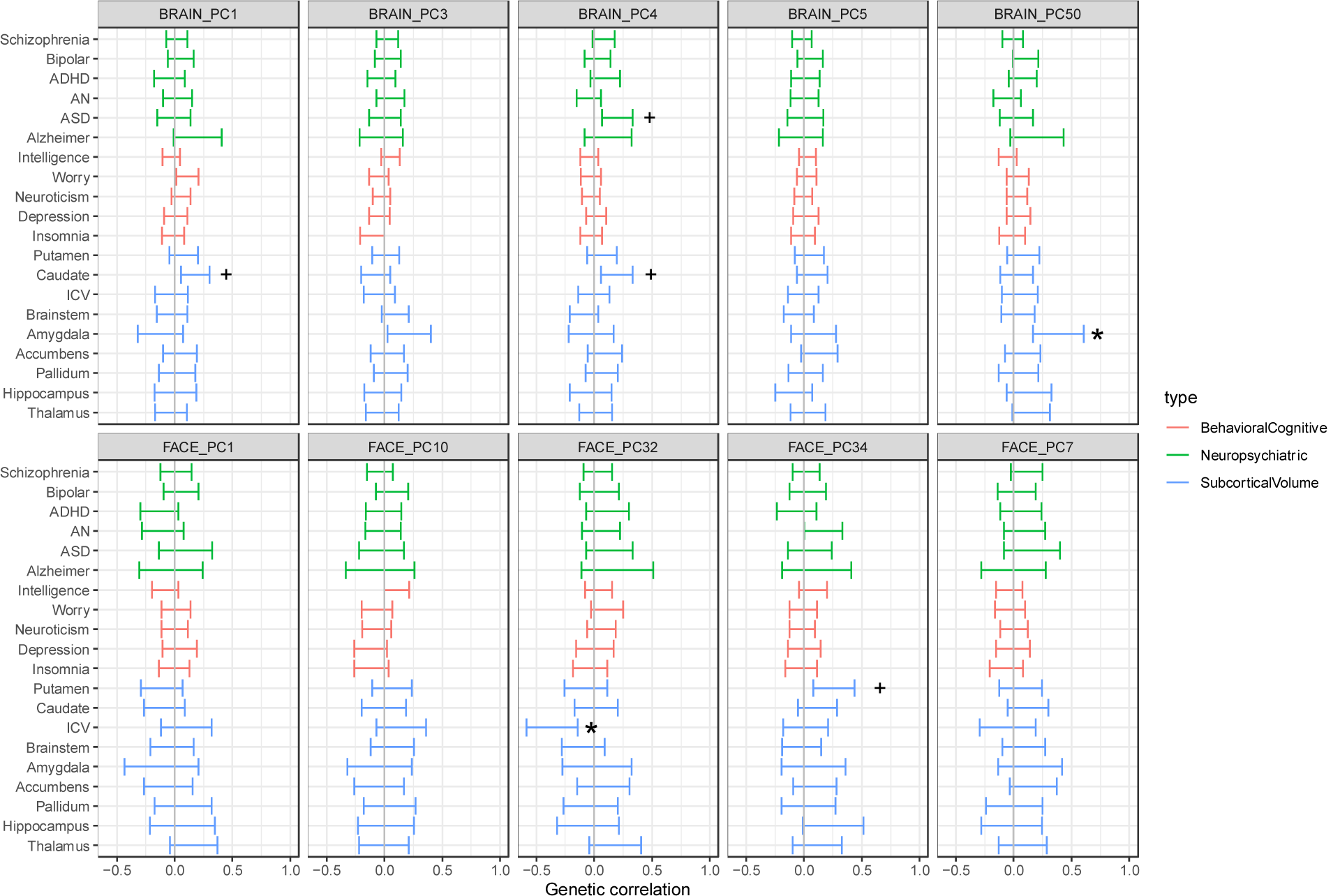
Genetic correlations between the most heritable brain (top) or face (bottom) shape PCs and other traits. Genetic correlations (rg) between the top five shape PCs (for segment 1, the full brain or face) with heritability z-score > 4 and each of the indicated univariate traits using LD score regression. Error bars represent 95% confidence intervals. *, 5% FDR for indicated PC; +, 10% FDR.

**Extended Data Fig. 9.**
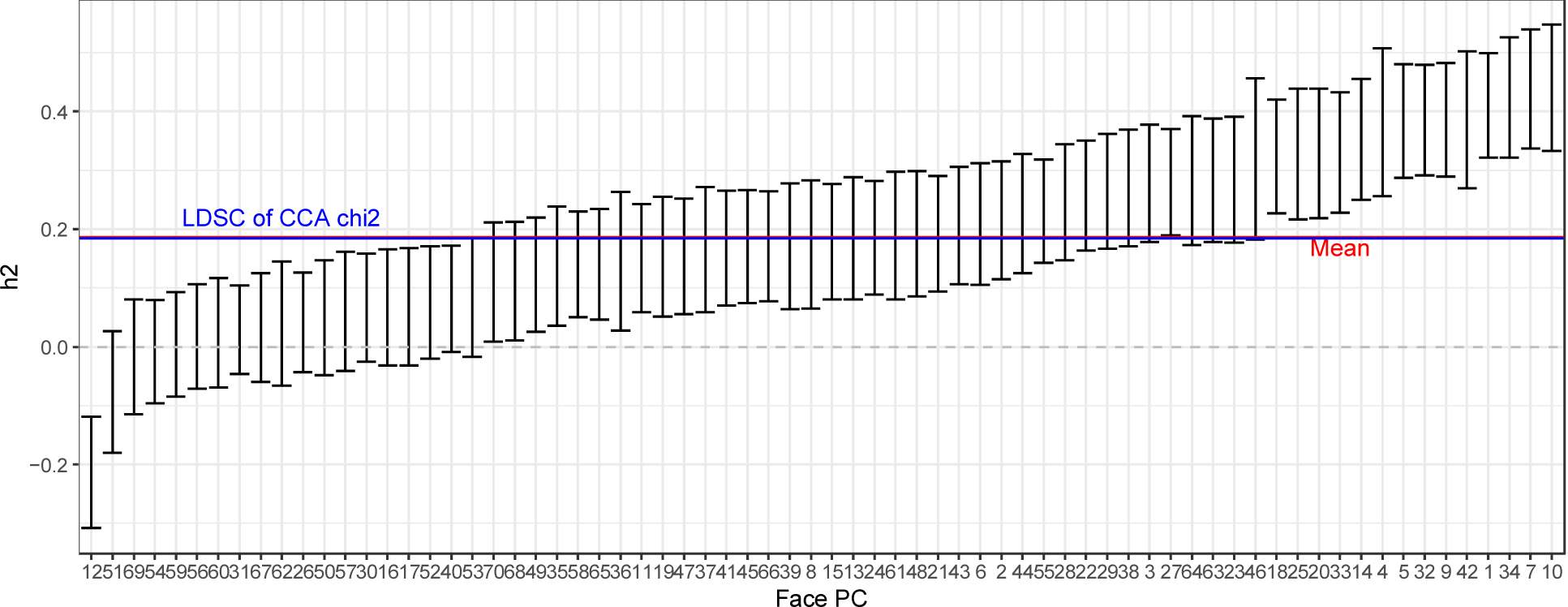
SNP heritability of individual face shape PCs and multivariate face shape estimated by LDSC. Error bars represent 95% confidence intervals. The red line represents the mean heritability of all 70 PCs, and the blue line indicates the heritability obtained by applying LDSC to corrected χ^2^ statistics from the multivariate CCA GWAS using all 70 PCs.

**Extended Data Fig. 10.**
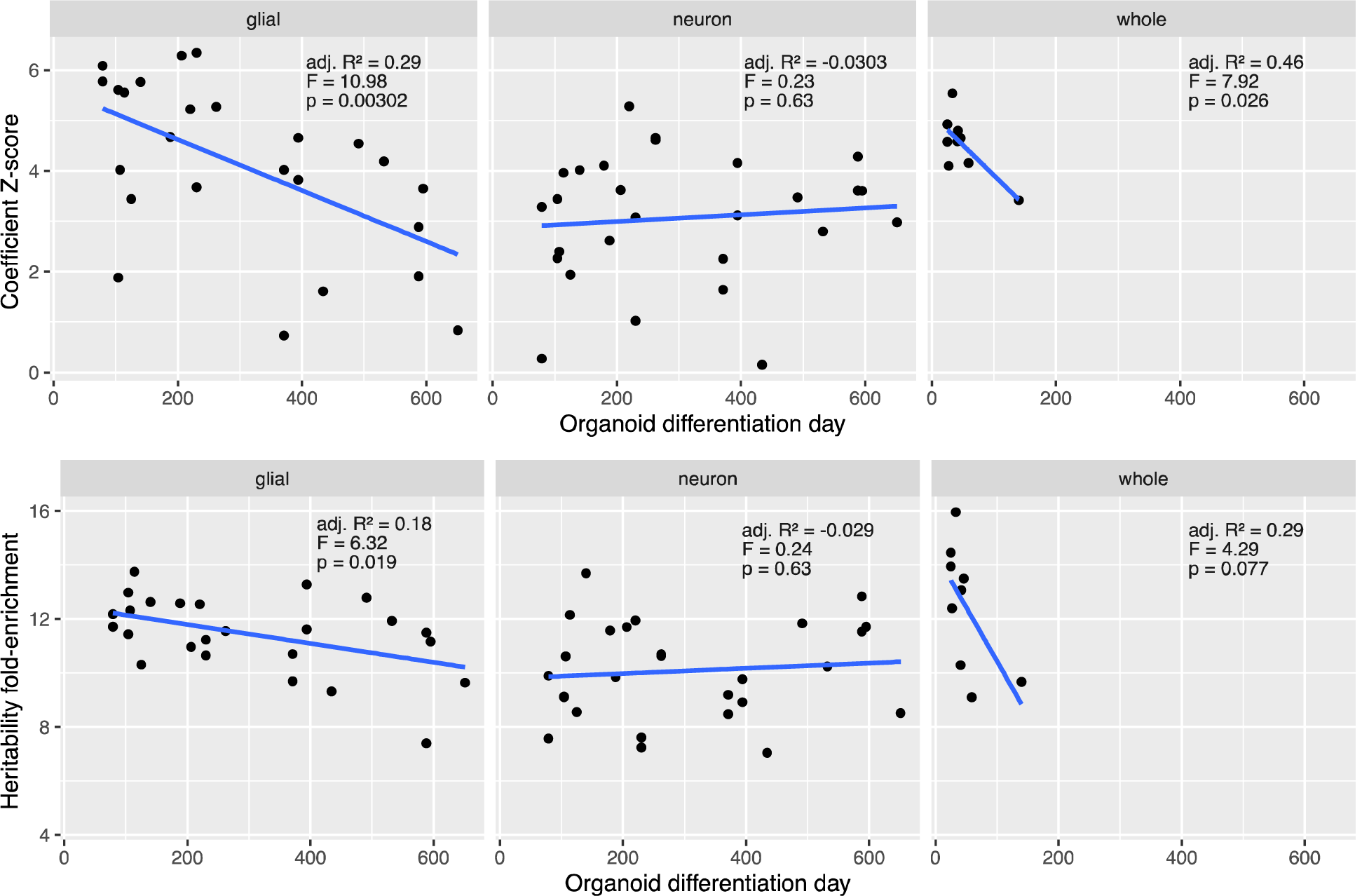
Partitioned heritability enrichments for brain shape with respect to stage- and cell-type-specific brain organoid open chromatin. S-LDSC coefficient Z-scores and heritability fold-enrichment for annotations corresponding to the indicated cell-type and differentiation day were computed as described in Methods. Regression lines represent the linear best fit with intercept and organoid differentiation day as dependent variable, and grey areas represent 95% confidence intervals.

**Extended Data Fig. 11.**
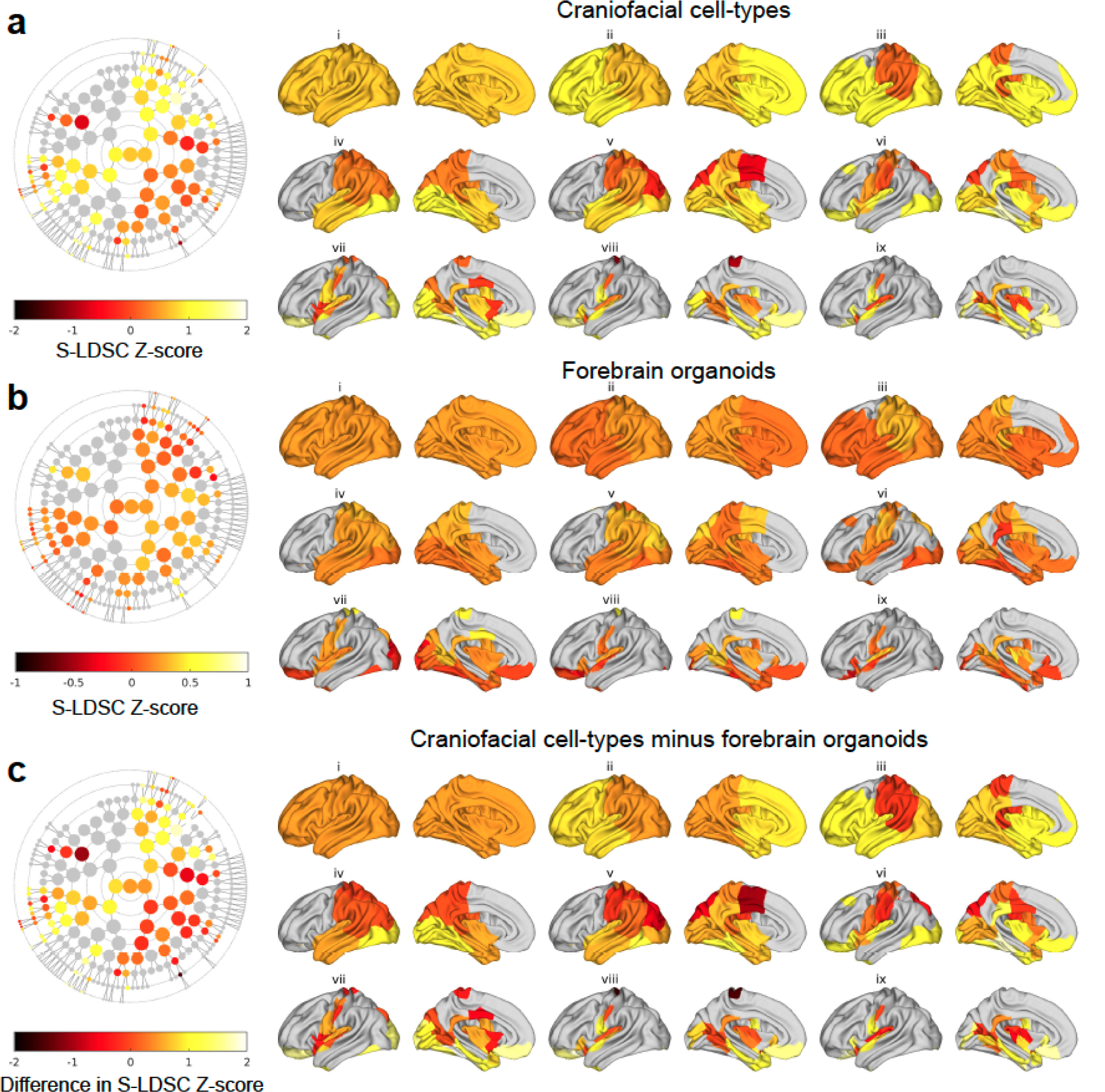
Partitioned heritability enrichments in craniofacial cell-types and brain organoids for brain shape within hierarchical segments. For each segment at the indicated hierarchical level, scaled S-LDSC Z-scores from all craniofacial (**a)** or brain organoid (**b)** annotations as indicated in Figure 4 were averaged, and the difference in mean Z-score between the two averages (**c**) was also computed.

**Extended Data Fig. 12.**
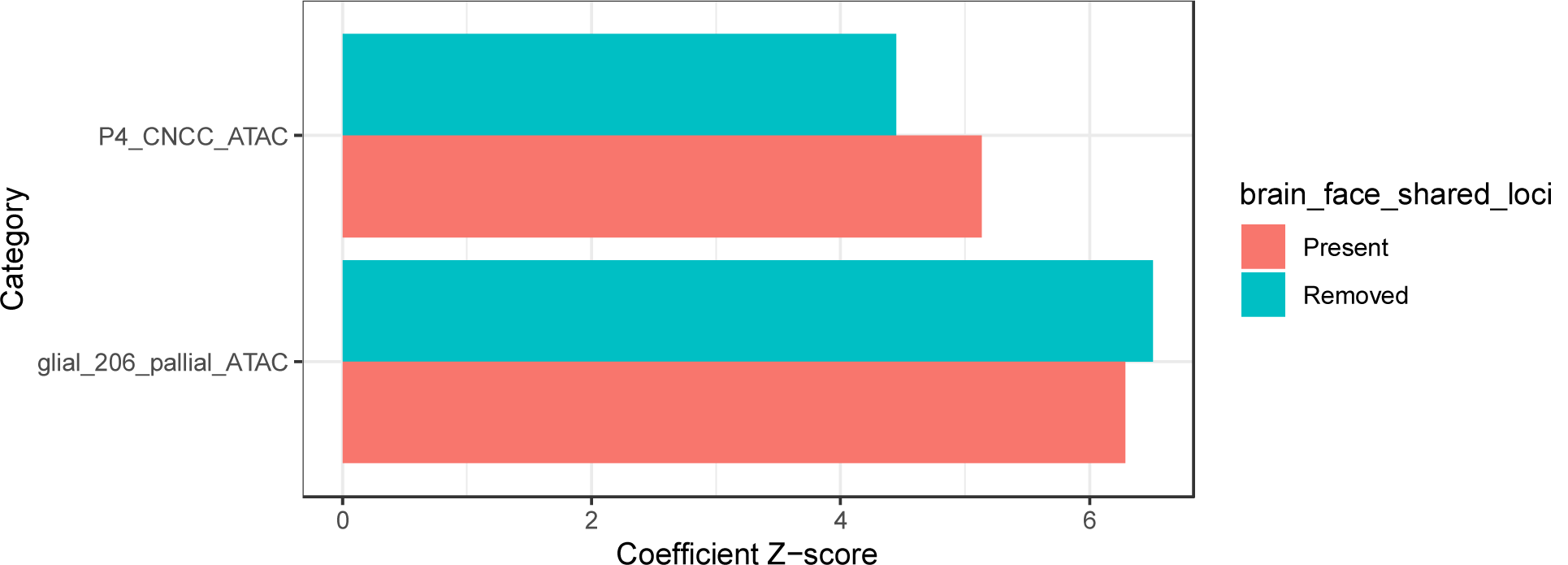
Partitioned heritability enrichments for brain shape with respect to open chromatin in CNCCs or early glial organoid cells, with or without 76 brain-face shared loci. S-LDSC Z-scores were calculated using full brain shape as the trait and the most enriched craniofacial (top) or brain organoid (bottom) ATAC-seq dataset as annotations. Z-scores were re-estimated (blue) after removing all SNPs in the same approximately independent LD block as one of the 76 brain-face shared loci (see Methods for details).

