## Supplementary material for "Shared heritability of face and brain shape distinct from cognitive traits": SI Guide

**Supplementary Table 1 Information on hierarchical segmentation of the mid-cortical surface.**
For each of the 285 hierarchical segments, ‘Level’ represents the hierarchical level of each segment, while ‘Cluster’ represents the cluster that segment was assigned to within that hierarchical level. These two variables can be used to look up the exact [x,y,z] coordinates of each segment’s vertices in the files provided on FigShare. The last four columns refer to the heritability of the segment as estimated by LDSC (see Methods, Supplementary Note)

**Supplementary Table 2 Information on 472 brain shape loci with genome-wide significant association with brain shape in at least one segment.**

The final four columns represent, for each SNP, the number of segments in each of the four quadrants (segments at hierarchical level 3) in which that SNP reached genome-wide significant association.

**Supplementary Table 3 Details on replication of 472 loci in the ABCD cohort.**

‘Segments2Test’ indicates the number of segments which that SNP showed genome-wide significant association in the discovery cohort, and which were tested for replication. ‘SNPTested’ indicates if the exact SNP was tested for replication (2), a proxy SNP was tested (1), or if that SNP was not tested (blank). ‘FDR_DEP’ and ‘Nominal’ indicate the number of tested segments that passed a 5% FDR threshold or a *P* < 0.05 cutoff, respectively.

**Supplementary Table 4 Summary of results from FUMA, GREAT, and gene panel enrichment analysis of the 472 brain shape loci.**

Excel file organized into 10 sheets, with each sheet containing significant terms by FUMA, GREAT, or gene panel analyses at three different distance cutoffs. A summary sheet with hierarchical relationships of tested gene panels, along with indication of significant panels, is also provided.

**Supplementary Table 5 Annotation of 76 brain-face shared loci.**

For each of the lead SNPs at the 76 brain-face shared loci (‘SNP’, see Methods for details), the -log10(p-value) for both the exact and proxy SNP in the brain or face shape GWAS is shown. Candidate genes at each locus (see Methods for details) are provided, along with notes indicating relevant functions annotated to each gene through gene ontology and literature searches.

**Supplementary Table 6 Spearman correlations between pairs of traits.**

For each pair of traits, the Spearman correlation statistic as well as bootstrapped p-value (see Methods) is provided. See Supplementary Table 1 for brain segment IDs used.

**Supplementary Table 7 Information on publicly available GWAS summary statistics used.**

**Supplementary Table 8 S-LDSC heritability enrichments.**

Each row represents the S-LDSC output for the indicated trait and annotation (‘Category’) added to the S-LDSC baseline model. ‘Segment’ refers to face or brain segment, NA otherwise.

**Supplementary Note: Proof the for application of LD score regression to multivariate trait GWAS**

Mathematical proof showing that when applied to corrected summary statistics from a multivariate GWAS, LDSC and S-LDSC provide the average heritability or heritability enrichment across the dimensions of the multivariate trait.
