## Supplementary Note for "Shared heritability of face and brain shape distinct from cognitive traits"

### Supplementary Note: LD Score Regression for Multivariate Traits

Jeffrey P. Spence

#### 1 Main Theoretical Results

In this note, we show that LDscore regression [1] may be used on the results of GWAS on multivariate traits, albeit with a slight difference in interpretation. In the following we consider a $D$ dimensional trait measured in a sample of $N$ individuals and assume that there are $P$ SNPs. We consider the standard additive model of phenotypes

$$\mathbf{Y} = \mathbf{X}\mathbf{B} + \mathbf{E} \tag{1}$$

where $\mathbf{Y} \in \mathbb{R}^{N \times D}$ is the matrix of phenotypes; $\mathbf{X} \in \mathbb{R}^{N \times P}$ is the standardized genotype matrix; $\mathbf{B} \in \mathbb{R}^{P \times D}$ is the matrix of effect sizes, with $\mathbf{B}_{pd}$ representing the effect of SNP $p$ on the $d^{\text{th}}$ dimension of the trait; and $\mathbf{E} \in \mathbb{R}^{N \times D}$ is the residual matrix, representing environmental or measurement noise. For notational convenience, we will use $\mathbf{x}_j$ to represent the $j^{\text{th}}$ column of the genotype matrix (i.e., the standardized genotypes of the $N$ individuals at SNP $j$).

The goal of this analysis is to use the p-values reported by a multivariate GWAS to learn something about the heritability of the trait. Before beginning, however, we note that for a univariate trait, under a purely additive model, heritability is defined as the proportion of phenotypic variance, $\sigma_P^2$ attributable to genetic variation $\sigma_G^2$. In the multivariate trait case, an appropriate definition is less clear because phenotypic variance is now represented by a $D \times D$ covariance matrix, $\Sigma_P$, and there is no single analogue of the proportion of variance explained by the genetic covariance matrix, $\Sigma_G$. The generalization that we will use is $\frac{1}{D}\text{trace}(\Sigma_G \Sigma_P^{-1})$, which reduces to the classical definition in the univariate case, and because $\Sigma_G \preceq \Sigma_P$, von Neumann's trace inequality shows that this number is bounded between 0 and 1, with the upper extreme being reached only when $\Sigma_G = \Sigma_P$. In some sense, this can be thought of as a coordinate system-free "average heritability" across the $D$ dimensions of the trait. Indeed, if either $\Sigma_G$ or $\Sigma_P$ are diagonal (i.e., the coordinates are either genetically uncorrelated or phenotypically uncorrelated respectively) then $\frac{1}{D}\text{trace}(\Sigma_G \Sigma_P^{-1})$ is exactly the average of the univariate heritabilities of each dimension. Furthermore, this definition of heritability is invariant to scaling the dimensions of the trait, for example by changing units of measurement. In Section 2 we show that this generalization of heritability is, in fact, the only definition of heritability that satisfies all of these properties.

In order to make analytical progress, we introduce some assumptions. We assume $N$ is large, and so we neglect terms of order $O(\frac{1}{N})$ that are induced by standardizing $\mathbf{Y}$ and $\mathbf{X}$. Like in the original derivation of LDscore regression [1] we assume that $\mathbf{X}$, $\mathbf{B}$, and $\mathbf{E}$ are all independent random variables with the following moment conditions

$$
\begin{aligned}
\mathbb{E}[\mathbf{X}] &= \mathbf{0}_{N\times P} \\
\mathbb{E}[\mathbf{B}] &= \mathbf{0}_{P\times D} \\
\mathbb{E}[\mathbf{E}] &= \mathbf{0}_{N\times D} \\
\mathbb{E}[\mathbf{X}_{ij}\mathbf{X}_{ik}] &= r_{jk} \\
\mathrm{Var}[(\mathbf{B}_{j1},\ldots,\mathbf{B}_{jD})] &= \frac{1}{P}\begin{pmatrix} h_1^2 & \rho_{12} & \cdots & \rho_{1D} \\ \rho_{12} & h_2^2 & & \\ \vdots & & \ddots & \\ \rho_{1D} & & & h_D^2 \end{pmatrix} \\
\mathrm{Var}[(\mathbf{E}_{i1},\ldots,\mathbf{E}_{iD})] &= \begin{pmatrix} 1-h_1^2 & \epsilon_{12} & \cdots & \epsilon_{1D} \\ \epsilon_{12} & 1-h_2^2 & & \\ \vdots & & \ddots & \\ \epsilon_{1D} & & & 1-h_D^2 \end{pmatrix}
\end{aligned}
$$

and we assume that the effect sizes at different SNPs are uncorrelated, and that the errors across individuals are uncorrelated (but violations of this assumption are discussed below). We make the further assumption that $\mathbf{Y}^T\mathbf{Y} = N\mathbf{I}_D$. This assumption says that the multivariate trait is defined such that the dimensions are uncorrelated in the sample, and that each dimension is standardized to have mean zero and unit variance. In Section 1.2 we relax this assumption to allow for dependent columns. We also assume that hypothesis testing is performed using the following test statistic for SNP $j$, with estimated effect sizes at that SNP, $\widehat{\mathbf{b}}_j^T := \frac{1}{N}\mathbf{x}_j^T\mathbf{Y}$:

$$
\chi_j^2 = N^2\widehat{\mathbf{b}}_j^T\left[(\mathbf{Y}-\mathbf{x}_j\widehat{\mathbf{b}}_j^T)^T(\mathbf{Y}-\mathbf{x}_j\widehat{\mathbf{b}}_j^T)\right]^{-1}\widehat{\mathbf{b}}_j, \tag{2}
$$

which, under our assumption of $\mathbf{Y}^T\mathbf{Y} = N\mathbf{I}_D$, we show that this statistic reduces to

$$
\chi_j^2 = \frac{N||\widehat{\mathbf{b}}_j||_2^2}{1-||\widehat{\mathbf{b}}_j||_2^2}
$$

and in the univariate case is

$$
\chi_j^2 = \frac{N\hat{\beta}_j^2}{1-\hat{\beta}_j^2}.
$$

This assumption is mostly for convenience – in Section 1.3 we show that this is essentially equivalent to Wilks' Lambda, the default test statistic in many software packages [6]. For large $N$ and $N \gg D$, the $\chi_j^2$ statistic is approximately $\chi^2$ distributed with $D$ degrees of freedom. As a result, a p-value, $p$, from a GWAS may be transformed into the $\chi_j^2$ statistic by taking the $p^{\text{th}}$ upper quantile of the $\chi^2$ distribution with $D$ degrees of freedom.

We note that this statistic is defined differently from that in the original LDscore regression papers [1, 4] even in the case of a univariate trait, where they use

$$
\chi^2_{\text{LDSC}} = N\hat{\beta}_j^2.
$$

The statistic in Equation 2 matches what is used in standard regression packages, whereas the statistic in the LDscore regression papers does not. When converting p-values to $\chi^2$ statistics for LDscore regression, care should be taken that the result is the formula in Equation 2, *not* the statistic used in LDscore regression as defined in [1].

To avoid this confusion, we define the $\chi_j^2$ statistic as would be produced by any standard regression package, and then frame our results in terms of this statistic.

**Theorem 1.** *Let the LD Score of SNP $j$ be defined as*

$$\ell_j := \sum_{k=1}^{P} r_{jk}^2.$$

*Then, with the assumptions described above, we have*

$$\mathbb{E}\left[\frac{\chi_j^2}{D\left(1+\frac{\chi_j^2}{N}\right)}\right] = \frac{N-1}{P}\left(\frac{\sum_{d=1}^{D} h_d^2}{D}\right)\ell_j + 1 + O\left(\frac{1}{N}\right). \tag{3}$$

*Proof.* First, note that under the assumption that $\mathbf{Y}^T\mathbf{Y} = N\mathbf{I}$ we have

$$\begin{aligned}
\chi_j^2 &= N^2\widehat{\mathbf{b}}_j^T\left[(\mathbf{Y}-\mathbf{x}_j\widehat{\mathbf{b}}_j^T)^T(\mathbf{Y}-\mathbf{x}_j\widehat{\mathbf{b}}_j^T)\right]^{-1}\widehat{\mathbf{b}}_j \\
&= N^2\widehat{\mathbf{b}}_j^T\left[\mathbf{Y}^T\mathbf{Y} - \mathbf{Y}^T\mathbf{x}_j\widehat{\mathbf{b}}_j^T - \widehat{\mathbf{b}}_j\mathbf{x}_j^T\mathbf{Y} + \widehat{\mathbf{b}}_j\mathbf{x}_j^T\mathbf{x}_j\widehat{\mathbf{b}}_j^T\right]^{-1}\widehat{\mathbf{b}}_j \\
&= N^2\widehat{\mathbf{b}}_j^T\left[N\mathbf{I}_D - N\widehat{\mathbf{b}}_j\widehat{\mathbf{b}}_j^T\right]^{-1}\widehat{\mathbf{b}}_j \\
&= \frac{N||\widehat{\mathbf{b}}_j||_2^2}{1-||\widehat{\mathbf{b}}_j||_2^2}
\end{aligned}$$

where the third equality follows from the definition of $\widehat{\mathbf{b}}_j$, and the final equality follows from the Sherman-Morrison formula.

This then directly implies that

$$\frac{\chi_j^2}{D(1+\frac{\chi_j^2}{N})} = \frac{N}{D}||\widehat{\mathbf{b}}_j||_2^2.$$

We therefore seek to compute the expected value of $||\widehat{\mathbf{b}}_j||_2^2$, with

$$\begin{aligned}
\mathbb{E}\left[||\widehat{\mathbf{b}}_j||_2^2\right] &= \mathbb{E}\left[\widehat{\mathbf{b}}_j^T\widehat{\mathbf{b}}_j\right] \\
&= \frac{1}{N^2}\mathbb{E}\left[\mathbf{x}_j^T\mathbf{Y}\mathbf{Y}^T\mathbf{x}_j\right] \\
&= \frac{1}{N^2}\mathbb{E}\left[\mathbf{x}_j^T\left(\mathbf{X}\mathbf{B}+\mathbf{E}\right)\left(\mathbf{X}\mathbf{B}+\mathbf{E}\right)^T\mathbf{x}_j\right] \\
&= \frac{1}{N^2}\Big(\mathbb{E}\left[\mathbf{x}_j^T\mathbf{X}\mathbf{B}\mathbf{B}^T\mathbf{X}^T\mathbf{x}_j\right] + \mathbb{E}\left[\mathbf{x}_j^T\mathbf{X}\mathbf{B}\mathbf{E}^T\mathbf{x}_j\right] \\
&\qquad + \mathbb{E}\left[\mathbf{x}_j^T\mathbf{E}\mathbf{B}^T\mathbf{X}^T\mathbf{x}_j\right] + \mathbb{E}\left[\mathbf{x}_j^T\mathbf{E}\mathbf{E}^T\mathbf{x}_j\right]\Big).
\end{aligned}$$

But, by the independence of $\mathbf{B}$, $\mathbf{E}$ and $\mathbf{X}$ and the moment condition $\mathbb{E}[\mathbf{E}] = \mathbf{0}_{N\times D}$ we have

$$\mathbb{E}\left[\mathbf{x}_j^T\mathbf{X}\mathbf{B}\mathbf{E}^T\mathbf{x}_j\right] = \mathbb{E}\left[\mathbf{x}_j^T\mathbf{E}\mathbf{B}^T\mathbf{X}^T\mathbf{x}_j\right] = 0.$$

Hence,

$$\mathbb{E}\left[||\widehat{\mathbf{b}}_j||_2^2\right] = \frac{1}{N^2}\Big(\mathbb{E}\left[\mathbf{x}_j^T\mathbf{X}\mathbf{B}\mathbf{B}^T\mathbf{X}^T\mathbf{x}_j\right] + \mathbb{E}\left[\mathbf{x}_j^T\mathbf{E}\mathbf{E}^T\mathbf{x}_j\right]\Big).$$

Tackling the first term on the right hand side, we have

$$\begin{aligned}\mathbb{E}\left[\mathbf{x}_j^T\mathbf{X}\mathbf{B}\mathbf{B}^T\mathbf{X}^T\mathbf{x}_j\right] &= \mathbb{E}\mathbb{E}\left[\mathbf{x}_j^T\mathbf{X}\mathbf{B}\mathbf{B}^T\mathbf{X}^T\mathbf{x}_j\middle|\mathbf{X}\right] \\ &= \mathbb{E}\left[\mathbf{x}_j^T\mathbf{X}\mathbb{E}\left[\mathbf{B}\mathbf{B}^T\middle|\mathbf{X}\right]\mathbf{X}^T\mathbf{x}_j\right] \\ &= \mathbb{E}\left[\mathbf{x}_j^T\mathbf{X}\mathbb{E}\left[\mathbf{B}\mathbf{B}^T\right]\mathbf{X}^T\mathbf{x}_j\right] \\ &= \frac{1}{P}\left(\sum_{d=1}^D h_d^2\right)\mathbb{E}\left[\mathbf{x}_j^T\mathbf{X}\mathbf{X}^T\mathbf{x}_j\right] \\ &= \frac{N^2}{P}\left(\sum_{d=1}^D h_d^2\right)\left(\ell_j + \frac{P-\ell_j}{N}\right) + O(1)\end{aligned}$$

where we used that $\mathbb{E}[\mathbf{B}\mathbf{B}^T]_{jj'} = \sum_{d=1}^D \mathbb{E}\mathbf{B}_{jd}\mathbf{B}_{j'd} = \sum_{d=1}^D \frac{h_d^2}{P}$ if $j = j'$ and 0 otherwise by the fact that effect sizes at different loci are uncorrelated and that $\mathbb{E}[\mathbf{x}_j^T\mathbf{X}\mathbf{X}^T\mathbf{x}_j] = N^2(\ell_j + \frac{P-\ell_j}{N}) + O(1)$ which was shown in [1].

Meanwhile,

$$\begin{aligned}\mathbb{E}\left[\mathbf{x}_j^T\mathbf{E}\mathbf{E}^T\mathbf{x}_j\right] &= \mathbb{E}\mathbb{E}\left[\mathbf{x}_j^T\mathbf{E}\mathbf{E}^T\mathbf{x}_j\middle|\mathbf{X}\right] \\ &= \mathbb{E}\left[\mathbf{x}_j^T\mathbb{E}\left[\mathbf{E}\mathbf{E}^T\middle|\mathbf{X}\right]\mathbf{x}_j\right] \\ &= \mathbb{E}\left[\mathbf{x}_j^T\mathbb{E}\left[\mathbf{E}\mathbf{E}^T\right]\mathbf{x}_j\right] \\ &= \left(\sum_{d=1}^D 1 - h_d^2\right)\mathbb{E}\left[\mathbf{x}_j^T\mathbf{x}_j\right] \\ &= N\left(\sum_{d=1}^D 1 - h_d^2\right)\end{aligned}$$

where we used similar calculations to tackle $\mathbb{E}[\mathbf{E}\mathbf{E}^T]$ as we did to tackle $\mathbb{E}[\mathbf{B}\mathbf{B}^T]$, relying on the noise being uncorrelated across individuals.

Combining these results, we see

$$\mathbb{E}\left[||\widehat{\mathbf{b}}_j||_2^2\right] = \frac{1}{P}\left(\sum_{d=1}^D h_d^2\right)\left(1 - \frac{1}{N}\right)\ell_j + \frac{D}{N} + O\left(\frac{1}{N^2}\right),$$

which, after multiplying by $N$ and dividing by $D$, implies

$$\mathbb{E}\left[\frac{\chi_j^2}{D\left(1 + \frac{\chi_j^2}{N}\right)}\right] = \frac{N-1}{P}\left(\frac{\sum_{d=1}^D h_d^2}{D}\right)\ell_j + 1 + O\left(\frac{1}{N}\right).$$

□

The practical implication of this theorem is that by regressing LD scores against the transformed $\chi_j^2$ statistics from a multivariate GWAS, we are able to infer the average of the heritabilities of the dimensions of the trait.

It is clear from the proof that the assumption that errors are uncorrelated across individuals is not strictly necessary: like LDscore regression, violations of this assumption would result in changes to the intercept term, but not to the slope. As such, test statistic inflation due to effects such as population structure will get captured by the intercept term without biasing the slope.

#### 1.1 Partitioning average heritability

While Theorem 1 assumes an infinitesimal model where each variant contributes equally to the average heritability, in reality there is reason to believe that SNPs with certain characteristics or SNPs in some regions of the genome may contribute more to heritability. For instance, SNPs that lie in open chromatin in relevant cell types might be expected to contribute more to heritability. In general, we may annotate each SNP as belonging to different categories, and we can then infer the average contribution to heritability of each distinct annotation, which may highlight which regions of the genome are important for the trait. These ideas were originally pioneered in the single dimension trait case in [4], and we extend those results to our present multidimensional trait case here.

To formalize this model, we partition the $P$ SNPs into non-overlapping annotations, and we call this partitioning $\mathcal{C}$. Let $P(c)$ denote the number of SNPs in partition $c$, so that $\sum_{c\in\mathcal{C}} P(c) = P$. We maintain the same moment conditions as before, except now we have that for each annotation $c \in \mathcal{C}$, for each SNP $j \in c$,

$$\mathrm{Var}\left[(\mathbf{B}_{j1}, \ldots, \mathbf{B}_{jD})\right] = \frac{1}{P(c)} \begin{pmatrix} h_1^2(c) & \rho_{12}(c) & \cdots & \rho_{1D}(c) \\ \rho_{12}(c) & h_2^2(c) & & \\ \vdots & & \ddots & \\ \rho_{1D}(c) & & & h_D^2(c) \end{pmatrix},$$

and we define the the heritability of a dimension as the sum of the heritabilities contributed from each annotation:

$$h_d^2 := \sum_{c\in\mathcal{C}} h_d^2(c).$$

That is, we allow the distribution of effect sizes for SNPs in each annotation to have an arbitrary variance-covariance matrix determined by that annotation.

With this generalization of the assumptions of Theorem 1 we obtain the following generalization.

**Theorem 2.** *Under the assumptions listed above, and letting the annotation-specific LD score of SNP j and annotation $c \in \mathcal{C}$ be defined as*

$$\ell(j, c) := \sum_{k\in c} r_{jk}^2$$

*we have*

$$\mathbb{E}\left[\frac{\chi_j^2}{D\left(1+\frac{\chi_j^2}{N}\right)}\right] = (N-1)\left[\sum_{c\in\mathcal{C}}\frac{\ell(j,c)}{P(c)}\times\frac{\sum_{d=1}^{D}h_d^2(c)}{D}\right]+1+O\left(\frac{1}{N}\right). \tag{4}$$

*Proof.* From the proof of Theorem 1, we have that

$$\mathbb{E}\left[\frac{\chi_j^2}{D\left(1+\frac{\chi_j^2}{N}\right)}\right] = \frac{N}{D}||\widehat{\mathbf{b}}_j||_2^2$$

and

$$||\widehat{\mathbf{b}}_j||_2^2 = \frac{1}{N^2}\Big(\mathbb{E}\left[\mathbf{x}_j^T\mathbf{X}\mathbf{B}\mathbf{B}^T\mathbf{X}^T\mathbf{x}_j\right]+\mathbb{E}\left[\mathbf{x}_j^T\mathbf{E}\mathbf{E}^T\mathbf{x}_j\right]\Big).$$

The first term on the right hand side will need to be recalculated because of the different moment condition on $\mathbf{B}$, but the second term remains unchanged. To begin, note that $\mathbb{E}\left[\mathbf{B}\mathbf{B}^T\right]_{jj'}=0$ if $j\neq j'$ by the independence of sites and the fact that $\mathbf{B}$ has mean zero. For the diagonal terms,

$$\begin{aligned}\mathbb{E}\left[\mathbf{B}\mathbf{B}^T\right]_{jj} &= \sum_{d=1}^{D}\mathbb{E}\mathbf{B}_{jd}^2\\ &= \frac{1}{P(c(j))}\sum_{d=1}^{D}h_d^2(c(j)),\end{aligned}$$

where we wrote $c(j)$ for the partition that contains SNP $j$.

Then, using the independence of $\mathbf{B}$ and $\mathbf{X}$, and defining $\hat{r}_{jk}:=\mathbf{x}_j^T\mathbf{x}_k/N$ we obtain

$$\begin{aligned}\mathbb{E}\left[\mathbf{x}_j^T\mathbf{X}\mathbf{B}\mathbf{B}^T\mathbf{X}^T\mathbf{x}_j\right] &= \mathbb{E}\left[\mathbf{x}_j^T\mathbf{X}\mathbb{E}\left[\mathbf{B}\mathbf{B}^T\right]\mathbf{X}^T\mathbf{x}_j\right]\\ &= N^2\sum_{c\in\mathcal{C}}\frac{\sum_{d=1}^{D}h_d^2(c)}{P(c)}\sum_{k\in c}\mathbb{E}\hat{r}_{jk}^2\\ &= N^2\left[\sum_{c\in\mathcal{C}}\left(\ell(j,c)+\frac{P(c)-\ell(j,c)}{N}\right)\frac{\sum_{d=1}^{D}h_d^2(c)}{P(c)}\right]+O(1)\\ &= (N^2-N)\left(\sum_{c\in\mathcal{C}}\frac{\ell(j,c)}{P(c)}\sum_{d=1}^{D}h_d^2(c)\right)+N\left(\sum_{d=1}^{D}h_d^2\right)+O(1),\end{aligned}$$

where the third equality follows from the fact that $\mathbb{E}\hat{r}_{jk}^2=r_{jk}^2+(1-r_{jk}^2)/N+O(1/N^2)$, which was proved in [1].

Recall that

$$\mathbb{E}\left[\mathbf{x}_j^T\mathbf{E}\mathbf{E}^T\mathbf{x}_j\right] = N\left(\sum_{d=1}^{D}1-h_d^2\right) = ND-N\sum_{d=1}^{D}h_d^2$$

so combining we have

$$||\widehat{\mathbf{b}}_j||_2^2 = \left(1-\frac{1}{N}\right)\left(\sum_{c\in\mathcal{C}}\frac{\ell(j,c)}{P(c)}\sum_{d=1}^{D}h_d^2(c)\right)+\frac{D}{N}+O\left(\frac{1}{N^2}\right)$$

which implies that

$$\mathbb{E}\left[\frac{\chi_j^2}{D\left(1+\frac{\chi_j^2}{N}\right)}\right] = (N-1)\left[\sum_{c\in\mathcal{C}}\frac{\ell(j,c)}{P(c)}\times\frac{\sum_{d=1}^{D} h_d^2(c)}{D}\right] + 1 + O\left(\frac{1}{N}\right).$$

□

An interesting consequence of this definition of heritability enrichment is that it is invariant to the combinations of dimensions that each annotation affects – all that matters is the average heritability of each dimension. As a concrete example, consider the following two models. In the first model, the annotations are meaningless and variants in each annotation have the same distribution of effects on the trait. In the second model, the SNPs in each annotation affect only a single dimension of the trait, and each annotation affects a different dimension of the trait, but the heritability of each dimension is the same. Under both of these models, the average heritability across dimensions of the trait is the same for each annotation, and so there is no enrichment of heritability in any annotation in either model.

#### 1.2 Traits with correlated dimensions

In the above, we assumed that $\mathbf{Y}$ was normalized and rotated such that $\mathbf{Y}^T\mathbf{Y} = N\mathbf{I}_D$. If $\mathbf{Y}$ is not normalized, we can perform the thin singular value decomposition of $\mathbf{Y} = \mathbf{U}\mathbf{S}\mathbf{V}^T$. Then, letting $\tilde{\mathbf{Y}} := \sqrt{N}\mathbf{U}$ we have $\tilde{\mathbf{Y}}^T\tilde{\mathbf{Y}} = N\mathbf{I}_D$ as required. We can then rewrite Equation 1 by noting that $\tilde{\mathbf{Y}} = \sqrt{N}\mathbf{Y}\mathbf{V}\mathbf{S}^{-1}$:

$$\tilde{\mathbf{Y}} = \mathbf{X}\tilde{\mathbf{B}} + \tilde{\mathbf{E}},$$

where $\tilde{\mathbf{B}} = \sqrt{N}\mathbf{B}\mathbf{V}\mathbf{S}^{-1}$ and $\tilde{\mathbf{E}} = \sqrt{N}\mathbf{E}\mathbf{V}\mathbf{S}^{-1}$. The proof of Theorem 1 requires us to compute $\mathbb{E}[\tilde{\mathbf{B}}\tilde{\mathbf{B}}^T]$ and $\mathbb{E}[\tilde{\mathbf{E}}\tilde{\mathbf{E}}^T]$. Below, we will write $\mathbf{\Sigma}_P = \frac{1}{N}\mathbf{Y}^T\mathbf{Y}$ for the phenotypic variation and $\mathbf{\Sigma}_G = P\text{Var}(\mathbf{B}_j)$ for the total genetic variance.

Beginning with $\mathbb{E}[\tilde{\mathbf{B}}\tilde{\mathbf{B}}^T]$ we have

$$\begin{aligned}\mathbb{E}[\tilde{\mathbf{B}}\tilde{\mathbf{B}}^T] &= N\mathbb{E}[\mathbf{B}\mathbf{V}\mathbf{S}^{-2}\mathbf{V}^T\mathbf{B}^T] \\ &= \text{trace}\left(\text{Var}(\mathbf{B}_j)\left(\frac{1}{N}\mathbf{Y}^T\mathbf{Y}\right)^{-1}\right)\mathbf{I}_P \\ &= \frac{1}{P}\text{trace}\left(\mathbf{\Sigma}_G\mathbf{\Sigma}_P^{-1}\right)\mathbf{I}_P.\end{aligned}$$

A similar calculation results in

$$\mathbb{E}[\tilde{\mathbf{E}}\tilde{\mathbf{E}}^T] = \text{trace}\left(\text{Var}(\mathbf{E}_i)\mathbf{\Sigma}_P^{-1}\right)\mathbf{I}_N.$$

Yet, under our additive model the phenotypic variation must equal the variance from the noise plus the genetic variance, and so we must have $\text{Var}(\mathbf{E}_i) = \Sigma_P - \Sigma_G$, ignoring terms of $O(1/N)$. Therefore,

$$\mathbb{E}[\tilde{\mathbf{E}}\tilde{\mathbf{E}}^T] = \mathbf{I}_N - \text{trace}\left(\mathbf{\Sigma}_G\mathbf{\Sigma}_P^{-1}\right)\mathbf{I}_N.$$

Using these results in the proofs of Theorems 1 results in the following generalization.

**Corollary.** *In the notation of Section 1.2, for general* $\frac{1}{N}\mathbf{Y}^T\mathbf{Y} = \mathbf{\Sigma}_P$*, we have*

$$\mathbb{E}\left[\frac{\chi_j^2}{1+\frac{\chi_j^2}{N}}\right] = \frac{N-1}{P}\text{trace}\left(\mathbf{\Sigma}_G\mathbf{\Sigma}_P^{-1}\right)\ell_j + D + O\left(\frac{1}{N}\right).$$

If we allow the distribution of effect sizes to change across genomic annotations like in Section 1.1, then we can consider the total contribution to genetic covariance of each annotation:

$$\mathbf{\Sigma}_G^c := P(c)\text{Var}\left(\mathbf{B}_j\right),$$

for any SNP $j \in c$, and we can let $\mathbf{\Sigma}_G := \sum_{c\in\mathcal{C}} \mathbf{\Sigma}_G^c$ denote the total contribution to genetic covariance across all SNPs. Following similar reasoning as above about $\mathbb{E}\left[\tilde{\mathbf{B}}\tilde{\mathbf{B}}^T\right]$ and $\mathbb{E}\left[\tilde{\mathbf{E}}\tilde{\mathbf{E}}^T\right]$ we see that we can simply replace $\sum_{d=1}^D h_d^2(c)$ by trace $\left(\mathbf{\Sigma}_G^c\mathbf{\Sigma}_P^{-1}\right)$ in the proof of Theorem 2 to obtain the following generalization.

**Corollary.** *In the notation of Section 1.2, for general* $\frac{1}{N}\mathbf{Y}^T\mathbf{Y} = \mathbf{\Sigma}_P$ *and allowing the distribution of effect sizes to change across a set of annotations,* $\mathcal{C}$*, that partitions the SNPs, we have*

$$\mathbb{E}\left[\frac{\chi_j^2}{1+\frac{\chi_j^2}{N}}\right] = (N-1)\left[\sum_{c\in\mathcal{C}}\text{trace}\left(\mathbf{\Sigma}_G^c\mathbf{\Sigma}_P^{-1}\right)\frac{\ell(j,c)}{P(c)}\right] + D + O\left(\frac{1}{N}\right).$$

#### 1.3 Using p-values from tests based on other test statistics

In previous work on multivariate traits [3] an alternative test statistic to Equation 2 was used. In particular, testing was performed using Wilks' lambda which is the default option for canonical correlation analysis-based multivariate regression in many software packages [6]. Here, we show that the p-values generated by tests based on Wilks' lambda are approximately equivalent to those based on the test statistic in Equation 2. This section is largely a recap of classical results [5, 2].

Wilks' lambda arises in the general multivariate regression setting defined by Equation 1. In this setting we are interested in testing whether any of several null hypotheses is false. This is often referred to as an omnibus test. In particular, consider the following null hypothesis

$$H_0 : \mathbf{CBA} = \mathbf{D}$$

for matrices $\mathbf{C} \in \mathbb{R}^{Q\times P}$, $\mathbf{A} \in \mathbb{R}^{D\times D}$ and $\mathbf{D} \in \mathbb{R}^{Q\times D}$. We may then define the following matrices

$$\mathbf{S}_e := \mathbf{A}^T(\mathbf{Y} - \mathbf{X}\hat{\mathbf{B}})^T(\mathbf{Y} - \mathbf{X}\hat{\mathbf{B}})\mathbf{A}$$
$$\mathbf{S}_h := (\mathbf{C}\hat{\mathbf{B}}\mathbf{A} - \mathbf{D})^T(\mathbf{C}(\mathbf{X}^T\mathbf{X})^{-1}\mathbf{C}^T)^{-1}(\mathbf{C}\hat{\mathbf{B}}\mathbf{A} - \mathbf{D}).$$

Finally, letting $\lambda_1, \ldots, \lambda_K$ be the non-zero eigenvalues of $\mathbf{S}_e^{-1}\mathbf{S}_h$, Wilks' lambda is defined as

$$\Lambda_{\text{Wilks'}} := \prod_{k=1}^{K}\frac{1}{1+\lambda_k}.$$

To specialize to the present case, we note that for GWAS, the tests are actually run marginally, so the model is

$$\mathbf{Y} = \mathbf{x}_j \mathbf{b}_j^T + \mathbf{E},$$

and we test against the null hypothesis

$$H_0 : \mathbf{b}_j^T = \mathbf{0}_{1\times D}.$$

That means that in the above omnibus hypothesis setting we have that $\mathbf{C} = 1$, $\mathbf{A} = \mathbf{I}_D$, and $\mathbf{D} = \mathbf{0}_{1\times D}$. As a result,

$$\begin{aligned}\mathbf{S}_e &= (\mathbf{Y} - \mathbf{x}_j\widehat{\mathbf{b}}_j^T)^T(\mathbf{Y} - \mathbf{x}_j\widehat{\mathbf{b}}_j^T) \\ \mathbf{S}_h &= \widehat{\mathbf{b}}_j(\mathbf{x}_j^T\mathbf{x}_j)^{-1}\widehat{\mathbf{b}}_j^T = N\widehat{\mathbf{b}}_j\widehat{\mathbf{b}}_j^T.\end{aligned}$$

and so

$$\mathbf{S}_e^{-1}\mathbf{S}_h = N\left[(\mathbf{Y} - \mathbf{x}_j\widehat{\mathbf{b}}_j^T)^T(\mathbf{Y} - \mathbf{x}_j\widehat{\mathbf{b}}_j^T)\right]^{-1}\widehat{\mathbf{b}}_j\widehat{\mathbf{b}}_j^T.$$

This matrix is the product of a full rank matrix and a matrix with rank one, so it is rank one and has only one eigenvalue. Because the trace of a matrix is the sum of its eigenvalues, the sole eigenvalue of this matrix must be the trace. We may then use the cyclic property of trace to find

$$\begin{aligned}\text{trace}(\mathbf{S}_e^{-1}\mathbf{S}_h) &= N\text{trace}\left(\left[(\mathbf{Y} - \mathbf{x}_j\widehat{\mathbf{b}}_j^T)^T(\mathbf{Y} - \mathbf{x}_j\widehat{\mathbf{b}}_j^T)\right]^{-1}\widehat{\mathbf{b}}_j\widehat{\mathbf{b}}_j^T\right) \\ &= N\text{trace}\left(\widehat{\mathbf{b}}_j^T\left[(\mathbf{Y} - \mathbf{x}_j\widehat{\mathbf{b}}_j^T)^T(\mathbf{Y} - \mathbf{x}_j\widehat{\mathbf{b}}_j^T)\right]^{-1}\widehat{\mathbf{b}}_j\right) \\ &= N\widehat{\mathbf{b}}_j^T\left[(\mathbf{Y} - \mathbf{x}_j\widehat{\mathbf{b}}_j^T)^T(\mathbf{Y} - \mathbf{x}_j\widehat{\mathbf{b}}_j^T)\right]^{-1}\widehat{\mathbf{b}}_j.\end{aligned}$$

Therefore Wilks' lambda in this case is

$$\Lambda_{\text{Wilks'}} = \frac{1}{1 + N\widehat{\mathbf{b}}_j^T\left[(\mathbf{Y} - \mathbf{x}_j\widehat{\mathbf{b}}_j^T)^T(\mathbf{Y} - \mathbf{x}_j\widehat{\mathbf{b}}_j^T)\right]^{-1}\widehat{\mathbf{b}}_j},$$

and

$$\frac{1 - \Lambda_{\text{Wilks'}}}{\Lambda_{\text{Wilks'}}} = N\widehat{\mathbf{b}}_j^T\left[(\mathbf{Y} - \mathbf{x}_j\widehat{\mathbf{b}}_j^T)^T(\mathbf{Y} - \mathbf{x}_j\widehat{\mathbf{b}}_j^T)\right]^{-1}\widehat{\mathbf{b}}_j.$$

A classical result says that when $Q = 1$, as in our case,

$$\frac{1 - \Lambda_{\text{Wilks'}}}{\Lambda_{\text{Wilks'}}}\frac{N - D + 1}{D}$$

is $F$ distributed with $D$ and $N - D + 1$ degrees of freedom (e.g., Equation 8.19 in [2]). This implies that in the limit of large $N$,

$$(N - D + 1)\frac{1 - \Lambda_{\text{Wilks'}}}{\Lambda_{\text{Wilks'}}} \sim \chi^2 \text{ with } D \text{ degrees of freedom}$$

Therefore, compared to the statistic used in Theorem 1, $\chi_j^2$, we have

$$(N - D + 1)\frac{1 - \Lambda_{\text{Wilks'}}}{\Lambda_{\text{Wilks'}}} = \left(1 - \frac{D}{N} + \frac{1}{N}\right)\chi_j^2 \approx \chi_j^2,$$

and p-values in both tests are computed under asymptotically equivalent distributions.

#### 2 A multivariate generalization of heritability

Above we used $\frac{1}{D}\text{trace}\left(\boldsymbol{\Sigma}_G\boldsymbol{\Sigma}_P^{-1}\right)$ as a D-dimensional generalization of heritability for a genetic covariance matrix $\boldsymbol{\Sigma}_G$ and a phenotypic covariance matrix $\boldsymbol{\Sigma}_P$. There are many other sensible generalizations that also reduce to the univariate definition of heritability, for instance based on the Frobenius norm $||\cdot||_F$ or any operator norm $||\cdot||_{\text{op}}$. A few examples include:

- $\dfrac{||\boldsymbol{\Sigma}_G||_F}{||\boldsymbol{\Sigma}_P||_F}$
- $\dfrac{\text{trace}(\boldsymbol{\Sigma}_G)}{\text{trace}(\boldsymbol{\Sigma}_P)}$
- $\dfrac{||\boldsymbol{\Sigma}_G||_{\text{op}}}{||\boldsymbol{\Sigma}_P||_{\text{op}}}$
- $||\boldsymbol{\Sigma}_G\boldsymbol{\Sigma}_P^{-1}||_F$
- $||\boldsymbol{\Sigma}_G\boldsymbol{\Sigma}_P^{-1}||_{\text{op}}$
- $||\boldsymbol{\Sigma}_P^{-1/2}\boldsymbol{\Sigma}_G\boldsymbol{\Sigma}_P^{-1/2}||_F$
- $||\boldsymbol{\Sigma}_P^{-1/2}\boldsymbol{\Sigma}_G\boldsymbol{\Sigma}_P^{-1/2}||_{\text{op}}$

...

Indeed, because all univariate norms are proportional, for any matrix norm $||\cdot||_M$, any ratio of the form $\frac{||\boldsymbol{\Sigma}_G||_M}{||\boldsymbol{\Sigma}_P||_M}$, or any properly scaled matrix norm applied to $\boldsymbol{\Sigma}_G\boldsymbol{\Sigma}_P^{-1}$ or $\boldsymbol{\Sigma}_P^{-1/2}\boldsymbol{\Sigma}_G\boldsymbol{\Sigma}_P^{-1/2}$ will reduce to the univariate definition of heritability. Note that for positive semi-definite matrices, trace$(\cdot)$ is equal to the nuclear norm, and so trace$(\cdot)$ is also a norm on the relevant space. There are, of course, many other sensible ways to generalize heritability.

To provide some justification for our particular generalization, we list four simple properties that any generalization of heritability should possess and show that our generalization is the only measure that satisfies these four properties. We list these properties informally, as well as in formal mathematical statements about a heritability function, $h^2(\cdot,\cdot)$ that maps a genetic covariance matrix $\boldsymbol{\Sigma}_G$ and a phenotypic covariance matrix $\boldsymbol{\Sigma}_P$ to a scalar. Throughout when we say for all $\boldsymbol{\Sigma}_G$ and $\boldsymbol{\Sigma}_P$, we implicitly mean only such pairs that satisfy the constraints required under a non-degenrate additive genetic model: $0 \preceq \boldsymbol{\Sigma}_G \preceq \boldsymbol{\Sigma}_P$ and $0 \prec \boldsymbol{\Sigma}_P$. These constraints simply mean that the phenotypic variance is at least as great as the genetic variance for any combination of the dimensions of the trait, and the phenotypic variance of any combination of dimensions of the trait is strictly positive.

1. *Invariant to units of measurement*

   Any measure of heritability should be independent of the units with which the dimensions of the trait are measured. Mathematically, $h^2(\boldsymbol{\Sigma}_G, \boldsymbol{\Sigma}_P) = h^2(\mathbf{M}\boldsymbol{\Sigma}_G\mathbf{M}, \mathbf{M}\boldsymbol{\Sigma}_P\mathbf{M})$ for any diagonal matrix $\mathbf{M} \succ 0$, and for any $\boldsymbol{\Sigma}_P$, and $\boldsymbol{\Sigma}_G$.

2. *Coordinate-free*

   The way we choose to delineate the trait into different dimensions is arbitrary, especially with respect to genetic and phenotypic variance. Genetic variants or environmental effects may act to alter specific combinations of dimensions as opposed to single dimensions. As such, the particular coordinate system we use to represent the trait should not impact our measure of heritability. Mathematically, $h^2(\mathbf{\Sigma}_G, \mathbf{\Sigma}_P) = h^2(\mathbf{U}\mathbf{\Sigma}_G\mathbf{U}^T, \mathbf{U}\mathbf{\Sigma}_P\mathbf{U}^T)$ for any orthogonal matrix $\mathbf{U}$.

3. *Linear in* $\mathbf{\Sigma}_G$

   If the variance attributable to genetics doubles, we would want the heritability to double. Similarly, the heritability of the trait attributable to two sets of independent variants should be the sum of the heritability attributed to each set.. That is, $h^2(c\mathbf{\Sigma}_G, \mathbf{\Sigma}_P) = ch^2(\mathbf{\Sigma}_G, \mathbf{\Sigma}_P)$ and $h^2(\mathbf{\Sigma}_G^{(1)} + \mathbf{\Sigma}_G^{(2)}, \mathbf{\Sigma}_P) = h^2(\mathbf{\Sigma}_G^{(1)}, \mathbf{\Sigma}_P) + h^2(\mathbf{\Sigma}_G^{(2)}, \mathbf{\Sigma}_P)$, for any scalar $c$, and any $\mathbf{\Sigma}_G$, $\mathbf{\Sigma}_G^{(1)}$, $\mathbf{\Sigma}_G^{(2)}$, and $\mathbf{\Sigma}_P$ such that the resulting matrices still obey the positive semidefinite ordering listed above.

4. *Maximized when* $\mathbf{\Sigma}_G = \mathbf{\Sigma}_P$

   When the genetic variance matches the phenotypic variance along all combinations of dimensions, the heritability should be 1. That is $h^2(\mathbf{\Sigma}, \mathbf{\Sigma}) = 1$ for any $\mathbf{\Sigma}$.

**Theorem 3.** *Let $h^2$ be a function that maps a pair of matrices, $\mathbf{\Sigma}_G$, and $\mathbf{\Sigma}_P$ such that $0 \preceq \mathbf{\Sigma}_G \preceq \mathbf{\Sigma}_P$ and $0 \prec \mathbf{\Sigma}_P$ to a scalar. Furthermore, assume that $h^2$ satisfies properties 1-4 listed above. Then,*

$$h^2(\mathbf{\Sigma}_G, \mathbf{\Sigma}_P) = \frac{1}{D}\text{trace}\left(\mathbf{\Sigma}_G\mathbf{\Sigma}_P^{-1}\right).$$

*Proof.* To begin, we can use the spectral decomposition $\mathbf{\Sigma}_P = \mathbf{U}_P\mathbf{\Lambda}_P\mathbf{U}_P^T$ and properties 1 and 2 to obtain:

$$\begin{aligned} h^2(\mathbf{\Sigma}_G, \mathbf{\Sigma}_P) &= h^2(\mathbf{\Sigma}_G, \mathbf{U}_P\mathbf{\Lambda}_P\mathbf{U}_P^T) && (\textit{spectral decomposition}) \\ &= h^2(\mathbf{U}_P^T\mathbf{\Sigma}_G\mathbf{U}_P, \mathbf{\Lambda}_P) && (\textit{Property 2}) \\ &= h^2(\mathbf{\Lambda}_P^{-1/2}\mathbf{U}_P^T\mathbf{\Sigma}_G\mathbf{U}_P\mathbf{\Lambda}_P^{-1/2}, \mathbf{I}_D) && (\textit{Property 1}) \\ &= h^2(\mathbf{\Sigma}_P^{-1/2}\mathbf{\Sigma}_G\mathbf{\Sigma}_P^{-1/2}, \mathbf{I}_D) && (\textit{spectral decomposition}) \end{aligned}$$

Therefore, $h^2(\mathbf{\Sigma}_G, \mathbf{\Sigma}_P)$ is equivalent to a function $\tilde{h}^2(\mathbf{\Sigma}_P^{-1/2}\mathbf{\Sigma}_G\mathbf{\Sigma}_P^{-1/2})$, that operates on a single matrix $0 \preceq \mathbf{\Sigma}_P^{-1/2}\mathbf{\Sigma}_G\mathbf{\Sigma}_P^{-1/2} \preceq \mathbf{I}_D$. It is clear that $\tilde{h}^2$ is linear in its argument by applying Property 3 to $h^2$:

$$\tilde{h}^2(\mathbf{M}_1 + \mathbf{M}_2) = h^2(\mathbf{M}_1 + \mathbf{M}_2, \mathbf{I}_D) = h^2(\mathbf{M}_1, \mathbf{I}_D) + h^2(\mathbf{M}_2, \mathbf{I}_D) = \tilde{h}^2(\mathbf{M}_1) + \tilde{h}^2(\mathbf{M}_2)$$

and

$$\tilde{h}^2(c\mathbf{M}) = h^2(c\mathbf{M}, \mathbf{I}_D) = ch^2(\mathbf{M}, \mathbf{I}_D) = c\tilde{h}^2(\mathbf{M}).$$

Furthermore, by Property 2 $\tilde{h}^2$ is also invariant to multiplication of its argument on the left and right by any orthogonal matrix, $\mathbf{U}$, and its inverse:

$$\tilde{h}^2(\mathbf{U}\mathbf{M}\mathbf{U}^T) = h^2(\mathbf{U}\mathbf{M}\mathbf{U}^T, \mathbf{U}\mathbf{I}_D\mathbf{U}^T) = h^2(\mathbf{M}, \mathbf{I}_D) = \tilde{h}^2(\mathbf{M})$$

Hence, we may diagonalize the argument of $\tilde{h}^2$ via its spectral decomposition, $\mathbf{M} = \mathbf{U}_M\mathbf{\Lambda}_M\mathbf{U}_M^T$:

$$\tilde{h}^2(\mathbf{M}) = \tilde{h}^2(\mathbf{U}_M\mathbf{\Lambda}_M\mathbf{U}_M^T) = \tilde{h}^2(\mathbf{\Lambda}_M),$$

but $\mathbf{\Lambda}_M$ only contains the eigenvalues of $\mathbf{M}$, so $\tilde{h}^2$ is a function of only the eigenvalues of its argument. Furthermore, permutation matrices are orthogonal, and so by left and right multiplying the argument by a permutation matrix and its inverse, we see that $\tilde{h}^2$ is unchanged. Therefore, $\tilde{h}^2$ is a linear, permutation-invariant function of the eigenvalues of its argument. The only functions that satisfies these properties are proportional to trace$(\cdot)$. This implies that $h^2(\mathbf{\Sigma}_G, \mathbf{\Sigma}_P) = C\text{trace}(\mathbf{\Sigma}_P^{-1/2}\mathbf{\Sigma}_G\mathbf{\Sigma}_P^{-1/2})$ for some constant of proportionality $C$. We may rearrange this to $h^2(\mathbf{\Sigma}_G, \mathbf{\Sigma}_P) = C\text{trace}(\mathbf{\Sigma}_G\mathbf{\Sigma}_P^{-1})$ by the cyclic property of the trace. Finally, Property 4 shows that the constant of proportionality must be $1/D$:

$$1 = h^2(\mathbf{\Sigma}, \mathbf{\Sigma}) = C\text{trace}(\mathbf{\Sigma}\mathbf{\Sigma}^{-1}) = C\text{trace}(\mathbf{I}_D) = CD \implies C = \frac{1}{D}.$$

Therefore, $h^2(\mathbf{\Sigma}_G, \mathbf{\Sigma}_P)$ can only be $\frac{1}{D}\text{trace}\left(\mathbf{\Sigma}_G\mathbf{\Sigma}_P^{-1}\right)$. $\square$

[2] Chris Chatfield and Alexander Collins. *Introduction to multivariate analysis*, volume 1. CRC Press, 1981.

[3] Peter Claes, Jasmien Roosenboom, Julie D White, Tomek Swigut, Dzemila Sero, Jiarui Li, Myoung Keun Lee, Arslan Zaidi, Brooke C Mattern, Corey Liebowitz, et al. Genome-wide mapping of global-to-local genetic effects on human facial shape. *Nature genetics*, 50(3):414–423, 2018.

[4] Hilary K Finucane, Brendan Bulik-Sullivan, Alexander Gusev, Gosia Trynka, Yakir Reshef, Po-Ru Loh, Verneri Anttila, Han Xu, Chongzhi Zang, Kyle Farh, et al. Partitioning heritability by functional annotation using genome-wide association summary statistics. *Nature genetics*, 47(11):1228, 2015.

[5] J Gary Lutz and Tanya L Eckert. The relationship between canonical correlation analysis and multivariate multiple regression. *Educational and Psychological Measurement*, 54(3):666–675, 1994.

[6] Uwe Menzel. CCP: Significance tests for canonical correlation analysis (CCA). *R package version*, 1, 2009.
